# A *Penicillium rubens* platform strain for secondary metabolite production

**DOI:** 10.1101/2020.04.05.026286

**Authors:** Carsten Pohl, Fabiola Polli, Tabea Schütze, Annarita Viggiano, László Mózsik, Sascha Jung, Maaike de Vries, Roel A.L. Bovenberg, Vera Meyer, Arnold J.M. Driessen

**Affiliations:** Molecular Microbiology, Groningen Biomolecular Sciences and Biotechnology Institute, University of Groningen, Groningen, The Netherlands; Applied and Molecular Microbiology, Institute of Biotechnology, TU Berlin, Berlin, Germany; DSM Biotechnology Centre, Delft, The Netherlands; Synthetic Biology and Cell Engineering, Groningen Biomolecular Sciences and Biotechnology Institute, University of Groningen, Groningen, The Netherlands

**Keywords:** Platform strain, Natural products, Penicillium, Biosynthetic gene clusters, Decumbenone, Calbistrin, CRISPR/Cas9, RNP

## Abstract

We present a *Penicillium rubens* strain with an industrial background in which the four highly expressed biosynthetic gene clusters (BGC) required to produce penicillin, roquefortine, chrysogine and fungisporin were removed. This resulted in a minimal secondary metabolite background. Amino acid pools under steady-state growth conditions showed reduced levels of methionine and increased intracellular aromatic amino acids. Expression profiling of remaining BGC core genes and untargeted mass spectrometry did not identify products from uncharacterized BGCs. This platform strain was repurposed for expression of the recently identified polyketide calbistrin gene cluster and achieved high yields of decumbenone A, B and C. The penicillin BGC could be restored through *in vivo* assembly with eight DNA segments with short overlaps. Our study paves the way for fast combinatorial assembly and expression of biosynthetic pathways in a fungal strain with low endogenous secondary metabolite burden.

## Introduction

The fungal kingdom contains a massive reservoir of biosynthetic gene clusters (BGCs) encoding secondary metabolites, offering the discovery of potential new natural lead compounds for numerous applications like pharmaceutical drugs or food ingredients. However, successful follow-up of fungal genome mining studies^1,2^ for novel secondary metabolites is often hampered by difficulties in handling promising fungi and identifying the specific conditions required for BGC activation^3–5^. To overcome these difficulties inherent to natural producers, a common approach is the transfer of promising BGCs into a more tractable organism. Although reported to be successful in prokaryotes like actinomycetes^6,7^ and eukaryotes such as baker’s yeast^8–10^ and filamentous fungi^11,12^, yields of heterologous secondary metabolites can vary substantially from the natural producer. Moreover, the native BGC-related product fingerprint of the host organism is mainly left unchanged, which drains cellular resources into unwanted BGC products, complicates target compound purification and can even lead to unspecific enzymatic conversion of the target compound if the specificity of native BGC enzymes is low. Thus, we envisioned that a tractable fungal host with a low background of endogenous secondary metabolites simplifies detection of novel molecules in broth samples as well as downstream purification.

*Penicillium rubens* (former name: *Penicillium chrysogenum*) is an industrially relevant fungal cell factory primarily used for production of ß-lactam-derived antibiotics. With the growing interest in fungal natural product discovery, precise genetic engineering of filamentous fungi gaining momentum^13^ and synthetic biology tools being frequently utilized to recode BGCs for heterologous hosts^14,15^, we reasoned that the specialization of *P. rubens* into a penicillin cell factory might be favorable for further advances towards a platform strain for expression of any novel BGC^16^. Similar approaches have been undertaken in *Aspergillus nidulans*^17^ where a strain with low background expression of endogenous BGCs was used for heterologous expression of BGCs randomly present on a large plasmids^11^. Several decades of classical strain improvement (CSI) have led to accumulation of point mutations^18^ that resulted in strains optimized for high ß-lactam yield in large scale fermenters^19^ and low unwanted secondary metabolite production. The superior fermentation characteristics of such strains were successfully employed for the production of cephalosporins^20^ and, after deletion of the penicillin BGC, also for the heterologous polyketide pravastatin^21^. *P. rubens* research has led to a full genome sequence^22^ and a metabolic model^23^ which makes it attractive for future rational strain improvements. In addition, the efficiency of integrating multiple DNA fragments into *P. rubens* has been increased by utilizing split-marker approaches^24^ and the targetable nuclease Cas9^25^. However, direct *in vivo* recombination for fast construction of different BGC pathway combinations has not yet been demonstrated. Moreover, since the precursors for the biosynthesis of penicillins, α-aminoadipic acid, L-cysteine and L-valine originate from diverse anabolic routes, a careful elucidation of intracellular amino acid pools would be required to assess the impact of CSI on the flexibility of the metabolism to respond to high and low amino acid demands.

Here, we report on the construction of a *P. rubens* strain lacking four highly expressed secondary metabolite BGCs resulting in a near to complete secondary metabolite deficient metabolome under the cultivation conditions tested here. We performed genomic and transcriptome analysis, characterized its amino acid profile and demonstrated its suitability for efficient BGC recombination by reconstructing the Penicillin BGC (Pen-BGC) of 17kb by in vivo recombination with 8 DNA fragment with short (110 bp) overlapping flanks. Finally, we utilized this new platform strain for expression of the heterologous calbistrin BGC from *P. decumbens*, yielding decumbenone A, B and C in the culture supernantant. This study paves the way to utilize *P. rubens* for exploration and production of novel BGCs.

## Results and Discussion

### 4 NRPSs display robust expression during low growth rates

To prioritize secondary metabolite BGCs from the industrial strain *P. rubens* DS68530 (derived from *Penicillium rubens* Wisconsin 54-1255 via CSI and targeted gene deletion, see **Figure 2c**), for deletion, expression levels of 49 annotated^22^ BGC core enzymes (nonribosomal peptide synthetases (NRPS), polyketide synthases (PKS) and hybrid enzymes thereof) were extracted from 22 publicly and 4 in-house available transcriptome data sets (**Supplementary Information SI1**) and grouped into high and low penicillin production conditions (**Supplementary Information SI2**). Penicillin yield was primarily affected by strain lineage and supplementation with side-chain precursors for production of Penicillin V and G. The growth rate was controlled by carbon limitation to alleviate the glucose repression of the Pen-BGC - mediated by CreA^26^. 17 out of 49 (35%) core BGC enzymes were not expressed in at least 22 of the 26 conditions covered by the transcriptome data and were therefore considered silent. However, besides the well-expressed ACV-tripeptide forming NRPS *pcbAB* (*Pc21g21390*), genes coding for three further NRPSs showed expression across multiple strain backgrounds under glucose-limited growth rates at 0.05 h^-1^ and below. As displayed in (**Figure** 1), these are *Pc21g12630* - *chyA*, located in the chrysogine BGC^27,28^, *Pc21g15480* - *roqA* in the roquefortine cluster^29^ and the NRPS producing fungisporin^30^ (*Pc16g04690* – *hcpA*). Four other BGC core genes (**Supplementary Information SI1**) showed fluctuating and lower expression under some conditions analyzed here, including Pc21g05070 (*sorA*) and Pc21g05080 (*sorB*), coding for 2 PKSs required for sorbicillin biosynthesis^31^. However, the mutation L146F in the ketosynthase domain of SorA of *P. rubens* DS68530 was shown to result in a non-functional enzyme^32^. Similar to this observation, increased expression of Pc16g11480 (encoding a PKS termed PKS7) was observed when strains were grown in shake flasks. PKS7 carries a mutation (A952D) in a putative linker region between domains, which could potentially affect enzyme functionality. Pc18g00380, coding for a NRPS-like gene (NRPS-like7) was constantly expressed at 10 to 15% of the actin expression level without having acquired any mutations during CSI, and no known metabolite has been associated to this core enzyme of a BGC. The remaining BGC core genes showed only very little expression and might thus not contribute significantly to the metabolite fingerprint of *P. rubens* under the tested cultivation conditions.

**Figure 1:**
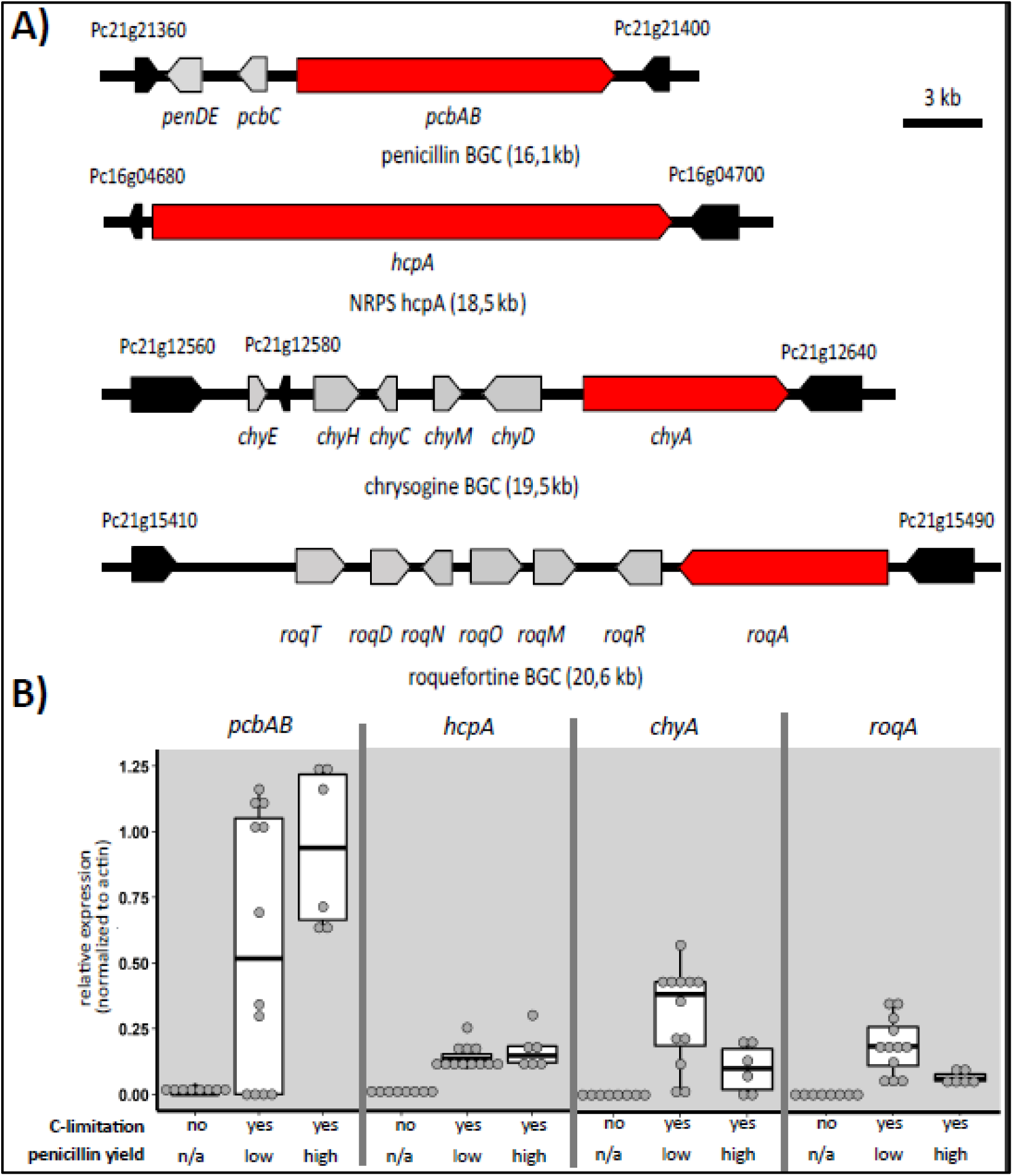
Genomic structure of four BGCs displaying strong expression and relative expression of the core gene under three different conditions. A) Schematic organization of the BGCs identified as being strongly expressed. BGC core genes are shown in red, closest genes not part of the cluster are colored black. All loci are drawn to scale and arrow directions denote orientation of transcription. B) Relative expression of identified BGC core genes from 26 transcriptome analyses with strong expression. Experiments were grouped according to carbon limitation and penicillin yield. n/a not applicable (because no quantities reported).

**Figure 2:**
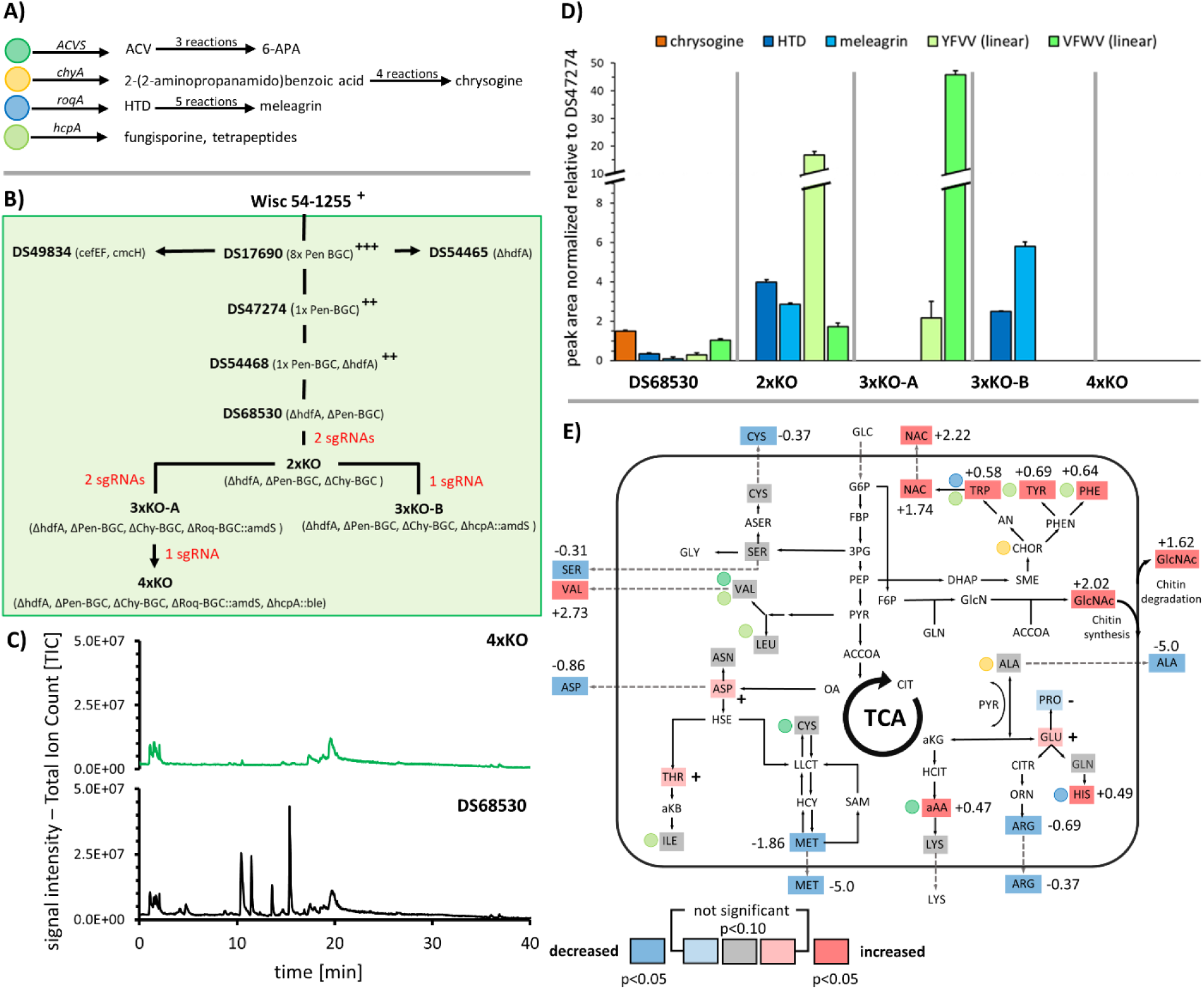
Effect of BGC deletion on secondary metabolites and amino acid levels. **A**) Overview of core BGC genes, first products in pathways and number of reactions steps leading to the final products 6-6-aminopenicillanic acid (6-APA), chrysogine, meleagrin and fungisporine. **B)** Strain lineage of *P. rubens*, including strains utilized in this study. Penicillin yields are denoted by superscript (+)-symbols ranging from (**+**) – low, (**++**) – intermediate to (**+++**)- high as far as reported. Figure adapted with modifications from^32^. **C)** Total-ion-chromatograms of DS68530 and 4xKO, taken after five days of growth in SMP. **D)** Changes in peak area of selected secondary metabolites associated with removed BGCs after five days of growth in SMP medium (n=3). A time-course series of all secondary metabolites can be found in **supplementary Information SI4**. E) Summary of changes in intra and extracellular amino acids and metabolites observed in the 4xKO strain compared to the penicillin-producing strain DS54468 cultivated at a growth rate of 0.05 in a glucose-limited chemostat. A schematic view of amino acid metabolism is shown. Values next to amino acids indicate log2 fold changes if significant. Decreases are indicated by blue, increases by red and unchanged amino acids by grey background. If the change is statistically not significant but p < 0.10, the increase or decline in concentration is denoted by (+) or (-), respectively. Abbreviations: NAC = nicotinic acid; HCIT = homocitrate; aAA = α-aminoadipic acid.

Overall, these data suggest that the identified BGCs have a stronger impact on the hosts secondary metabolite fingerprint due to increased product formation when the penicillin BGC is lower expressed or completely deleted. Indeed, the encoded NRPSs belong to BGCs which were characterized previously as abundant products in the culture broth^29,30,33^ of strains lacking the penicillin BGC. Therefore, the chrysogine and roquefortine BGCs along with the fungisporin-producing NRPS *hcpA* were prioritized for deletion.

### Construction and genomic analysis of a *P. rubens* strain devoid of four BGCs

A recently developed methodology of Cas9-aided transformation^25^ was used for sequential and complete deletion of the prioritized BGCs and the intermediate strains 2xKO (*ΔhdfA, Δpen-BGC, Δchy-BGC*), 3xKO-A (*ΔhdfA, Δpen-BGC, Δchy-BGC, Δroq-BGC::amdS*), 3xKO-B (*ΔhdfA, Δpen-BGC, Δchy-BGC, ΔhcpA::amdS*) and 4xKO (*ΔhdfA, Δpen-BGC, Δchy-BGC, Δroq-BGC::ergA, ΔhcpA::ble*) were obtained (**Figure 2 and Table 1**). Due to repeated transformations and treatment with Cas9 RNPs, the genome of *P. rubens* 4xKO was analyzed for mutations not previously reported in this strain lineage ^18^, using DS68530 as the reference strain. We observed 46 mutations in 4xKO, of which 18 mutations were located in 12 genes and the remaining mutations were intergenic. Remarkably, 13 of the 46 identified mutations were also present in DS68530 at frequencies below 50% in the population of spores used to generate material for gDNA and sequencing, suggesting that these mutations were then further enriched during clone selection and strain construction steps we conducted. The 18 genic mutations were classified as frameshift (1), intronic (1), non-synonymous SNP (12) or synonymous SNPs (4). Mutated genes encode for low-expressed hypothetical proteins where no clear biological role was immediately conclusive (**Supplementary Information SI3)** and did not contain a sgRNA off-target-site. Additionally, we also Sanger-verified the putative off-target sites with less than 4 mismatches and no bulges on the RNA or DNA of the five sgRNAs used during the transformation by PCR amplification and amplicon sequencing. None of these off-target sites did contain a mutation (**Supplementary Information SI4**), hence the applied Cas9-RNP method did not lead to off-target mutations in our hands, but the process of repeated transformations and selection caused a few novel mutations. This is not surprising as removal of the 8 pen-BGC copies from DS17690 also resulted in an accumulation of 18 previously not observed genic SNPs in the derived strain DS68530^34^. Moreover, it was also reported for *Aspergillus fumigatus*^35^ that spontaneous mutations can occur in the absence of Cas9. Also, no larger deletions or insertions were identified among the SNPs which were observed in *Aspergillus niger* capable of performing NHEJ^36^. The *P. rubens* strains utilized here are devoid of *hdfA*/*ku70* and thus, homology-directed repair (HDR) will be the dominant mechanism for DNA damage, therefore additionally limiting the possibility of mutations. Taken together, the obtained 4xKO strain does carry very few new mutations compared to the parental strain DS54468 which are not caused by repeated treatment with Cas9 RNPs, stressing that our applied methodology is efficient and reliable.

**Table 1:** Strains created in this study and transformations performed.

| Strain | genotype | parental strain | donor DNA strategy | clones total | µg marker cassette used | tested / positive clones (colony PCR) |
| --- | --- | --- | --- | --- | --- | --- |
| 2xKO | (ΔhdfA, ΔPen-BGC, ΔChy-BGC ) | DS68530 | 1 part, 1500 bp homology, marker free | 67 | 10 | 24/2 |
| 3xKO-A | (ΔhdfA, ΔPen-BGC, ΔChy-BGC, ΔRoq-BGC::amdS ) | 2xKO | 1 part, 100 bp homology, acetamide selection | 106 | 5 | 6/6 |
| 3xKO-B | (ΔhdfA, ΔPen-BGC, ΔChy-BGC, ΔhcpA::amdS ) | 2xKO | 1 part, 100 bp homology, acetamide selection | 213 | 5 | 6/6 |
| 4xKO | (ΔhdfA, ΔPen-BGC, ΔChy-BGC, ΔRoq-BGC::amdS, ΔhcpA::ble) | 3xKO-A | 1 part, 100 bp homology, phleomycin selection | 79 | 5 | 6/4 |
| 4xKO-B | (ΔhdfA, ΔPen-BGC, ΔChy-BGC, ΔRoq-BGC::ergA, ΔhcpA:: ble) | 4xKO | 1 part, 100 bp homology, terbinafine selection | 238 | 5 | 6/5 |
| **4xKO-B-PenBGC** | pen-BGC in IGR | 4xKO-B | 1 part, >1000bp<br>homology, acetamide<br>selection | 11 | 3 | 6/6 |
| **DS68530-PenBGC** | pen-BGC in IGR | DS68530 | 4 parts, >1000 bp<br>homology, acetamide<br>selection | 41 | 3 | 6/6 |
| **4xKO-B-PenBGC-p40s** | pen-BGC in IGR, p40s for<br>all genes | 4xKO-B | 8 parts, 100 bp<br>homology, acetamide<br>selection | 26 | 2 | 17/4 |
| **DS68530-**<br>**PKS17-OE** | Integrating pIPNS in front<br>of Pc21g16000 | DS68530 | 1 part, 100 bp<br>homology, acetamide<br>selection | 146 | 4 | 6/6 |
| **4xKO-B-**<br>**PKS17-OE** | Integrating pIPNS in front<br>of Pc21g16000 | 4xKO-B | 100 bp homology,<br>acetamide selection | 183 | 4 | 6/6 |
| **4xKO-B-Cal-BGC** | Integrating cal-BGC in IGR | 4xKO-B | 6 parts, 100 to >1000<br>bp homology,<br>acetamide selection | 97 | 3 | 16/16 |

### *P. rubens* devoid of four BGCs shows low secondary metabolite levels

The 4xKO strain as well as the intermediate 2xKO, 3xKO-A and 3xKO-B strains, and the parental strain DS47274 (1x *pen-BGC*) (see Figure 2b) were subjected to shake-flask cultivations using secondary metabolite production (SMP) medium^37^. The culture supernatant was subjected to LC-MS to identify changes in secondary metabolite profiles from day 2 to 7 of cultivation (**Figure 2, Supplementary Information SI5, Supplementary information SI6**).

Importantly and as expected, the total ion chromatograms of untargeted LC-MS runs (**Supplementary information SI7A**) of the 4xKO strain revealed that all penicillin, chrysogine, roquefortine and fungisporin-related metabolites were absent. In the intermediate strains, differential responses in secondary metabolite profiles were apparent. As shown in Figure 2, the level of chrysogine increased moderately (1.5 fold on day 5) upon deletion of the Pen-BGC while the level of modified chrysogines (chrysogine 6, 9 and 10) increased up to 29-fold (**Supplementary Information SI6**). However, these compounds are only a minor fraction (with normalized peak areas between 10^3^ to 10^5^) of all produced secondary metabolites whereas chrysogine is present in high quantities (normalized peak areas at 10^7^) already after 48 h but does not accumulate further throughout the cultivation (**Supplementary Information SI6**). Besides, the levels of histidyltryptophanyldiketopiperazine (HTD) and meleagrin decreased to 0.35- and 0.1-fold of wildtype levels, respectively, while fungisporin related compounds (e.g. YFVV, VFWV) remained almost unchanged.

After deletion of the chrysogine BGC (Chy-BGC), an almost 4-fold increase of HTD levels (normalized peak areas at 10^6^) and a 2.8-fold increase in meleagrin levels was observed together with up to 16-fold more of the linearized fungisporin^30^ tetrapeptides consisting of tyrosine, phenylalanine and valine (YFVV, VYFV, FVVY) while tryptophan-containing tetrapeptides (YWVV, VYWV, WVVY) increased only by 1.7-fold (**Figure 2c**). Upon deletion of the roquefortine BGC (Roq-BGC), the product pattern of fungisporins was changed and a 45-fold increase of tryptophan-containing tetrapeptides was detected, whereas the increase in non-tryptophan-containing tetrapeptides was only 2-fold. Similarly, when the fungisporin NRPS gene *hcpA*^30^ was deleted, a 5.8-fold increase of meleagrin compared to levels in DS27472 was observed. Additionally, expression of remaining BGC core genes in the 4xKO strain were quantified using qPCR after 5 days of growth in shake flasks using SMP (**Supplementary information SI8A)**. Of the 45 remaining SM core genes, 7 genes showed an increased expression with log2 FC > 2, however all genes were expressed with a relative expression level below 5% of the reference actin. The extracellular metabolome of the 4xKO strain was also analyzed for the appearance of novel peaks and filtered for changes in m/z abundance using XCMS^38^. Albeit we detected numerous m/z features with a log2 FC greater than 2, only 5 features had a relative peak area greater than 1×10E6. Also, multiple features with an m/z greater than 750 (**Supplementary information SI7**) were observed. Since no proteins were precipitated prior to analysis of the broth, it might be possible that some of these higher molecular weight features can be related to protein fragments. In conclusion, we did not observe unexpected changes in the secondary metabolites excreted by the 4xKO strain, which represents astrain with a reduced extracellular secondary metabolite metabolome we characterized further on.

### Characterization of *P. rubens* devoid of four BGCs using chemostat cultivation

LC-MS data indicated changed patterns in BGC related products of intermediate strains which are likely caused by altered amino acid availability due to deletion of BGCs that require specific amino acids and thus draining the available pools for these amino acids. Therefore, a quantification of intracellular amino acids levels in the 4xKO strain was performed under steady-state growth conditions, which can be controlled in glucose-limited chemostats.

The maximum growth rate (µ_max_) on glucose of two glucose-limited batch cultivations was found to be similar, i.e. 0.15 ± 0.001 h^-1^ and 0.14 ± 0.003 for DS54468 and 4xKO, respectively. A growth rate of 0.05 h^-1^ was selected for glucose-limited chemostat cultivations for both strains, as under these conditions there is acceptable production of penicillin^39^ while this resembles the growth rate of *P. rubens* on lactose^40,41^, the carbon source used in SMP medium. A comparison of biomass concentration at several points during steady state between both strains revealed a slight increase of 6% in biomass concentration (6.59 ± 0.153 g/kg for DS54468 and 6.98 ± 0.054 g/kg for 4xKO; p=0.0034, two-tailed students t-test, n=5 for DS54468 and n= 7 for 4xKO) while dissolved oxygen tension (DOT), CO_2_ production, O_2_ consumption and base addition remained unchanged (**Supplementary Information SI9**)). Morphology of both strains was regularly checked microscopically. Both strains appeared similar in length and aggregation of hyphae during exponential and steady state phase (**Supplementary Information SI10**).

To examine possible changes in amino acid pool in the 4xKO strain, mycelium samples from chemostat cultivations of DS54468 and 4xKO were analyzed using LC-MS. The analysis of intra- and extracellular amino acids (**Figure 2d**) indicated an overall modest change in amino acid levels with 13 out of 19 quantified intracellular amino acids remaining unchanged. However, a significant change occurred for sulfur-containing and aromatic amino acids. While intracellular levels of cysteine remained unaltered (0.8 ± 0.4 µM/ g CDW), the extracellular level decreased from 2.3 nM to 1.7 nM (−0.37 log2 FC) whereas the intracellular concentration of methionine was reduced from 0.09 ± 0.03 µM/ g CDW in DS54468 to 0.02 ± 0.01 µM/ g CDW in the 4xKO strain (log2 FC of −1.86) and was completely undetectable in the culture supernatant of 4xKO (DS54468: 0.2 nM, log2 FC of −5.0). All aromatic amino acids increased at a log2 FC of around 0.5 (Trp: 0.11 ± 0.05 µM/g CDW; Tyr: 0.42 ± 0.07 µM/g CDW; Phe: 0.55 ± 0.06 µM/g CDW) in the 4xKO strain. Additionally, an increase of intracellular (log2 FC of 1.74) and extracellular (log2 FC of 2.22) nicotinic acid (NAC, [m+H]^+^ = 124.0393 m/z) was observed. This metabolite is produced from tryptophan and acts as a precursor of nicotinamide adenine dinucleotide (NAD). The intracellular level of valine remained unchanged (3.2 ± 0.8 µM/g CDW), however the valine level in the culture broth increased from 0.03 nM to 0.23 nM (2.73 log2 FC) and the intracellular concentration of α-amino adipic acid increased by a log2 FC of 0.47 in the absence of the penicillin BGC. Similarly, a moderate increase of histidine, the precursor for HTD and roquefortine was measured with concentrations increasing from 1.2 ± 0.2 µM/g CDW in DS54468 to 1.7 ± 0.3 µM/ g CDW in the 4xKO strain (log2 FC of 0.49). While the intracellular level of alanine, a precursor for chrysogine, remained unchanged (32.5 ± 4.5 µM/ g CDW) no extracellular alanine was detected in the culture supernatant of 4xKO (DS54468: 6.5 nM, log2 FC of −5.0). The intracellular and extracellular level of GlcNAc increased by log2 FC 2.02 and 1.62, respectively. Overall the changes we observe here are modest but indicate the increased levels of amino acids in the absence of utilizing BGCs.

### Transcriptome profile of *P. rubens* 4xKO displays distinct expression changes

Samples for transcriptome analysis by RNA-sequencing (RNA-seq) were taken from steady state chemostat cultivations and analyzed for changes in transcripts, resulting in 4274 differently expressed genes (**Supplementary Information SI11 and SI12**). Out of the 45 remaining BGC core genes, three NRPS-like and a single PKS (Pc21g00960) showed a log2 FC >2, however, none of these genes was expressed above a level of 2% relative to actin, showing that silent BGC clusters were not activated in our 4xKO strain consistent with the lack of new secondary metabolites in the extracellular metabolite profile.

We next aimed to identify changes in transcript abundance that resulted from absence of the four BGCs and are not solely due to a reduction of the penicillin synthesis burden. We retrieved expression changes from available microarray datasets comparing high-versus-low penicillin production condition for the differently expressed genes identified by RNA-seq and calculated z-scores for each gene (**Figure 3a**). This was possible for 3834 genes (87.9 % of the differently expressed genes) with trackable expression behavior in at least 50% of the considered microarray datasets. Among these, 1594 genes (41.6%) were expressed very similar (|z| < 0.2) compared to strains where production pressure was reduced by omitting the addition of the penicillin sidechain precursors phenoxyacetic acid or phenylacetic acid to the culture medium. A subset of 2077 genes showed an altered expression with 0.2 < z < −0.2. Very different expression behavior was observed for 82 genes with z above 1.25 and 81 genes had a z-score below −1.25, mainly covering BGC-related genes that were deleted (**Figure 3b and Supplementary Information SI13**). It was also possible to track genes with a higher log2-FC showing similar responses (**Supplementary Information SI14**). FunCat enrichment analysis (**Figure 3c and Supplementary Information SI14**) of genes with |z| >0.2 identified 20 categories as enriched (p < 0.05), amongst them metabolism, biosynthesis and degradation of phenylalanine, supporting further analysis of the gene set by mapping it to KEGG^42^ Pathways (**Figure 3d and Supplementary Information SI16**).

**Figure 3:**
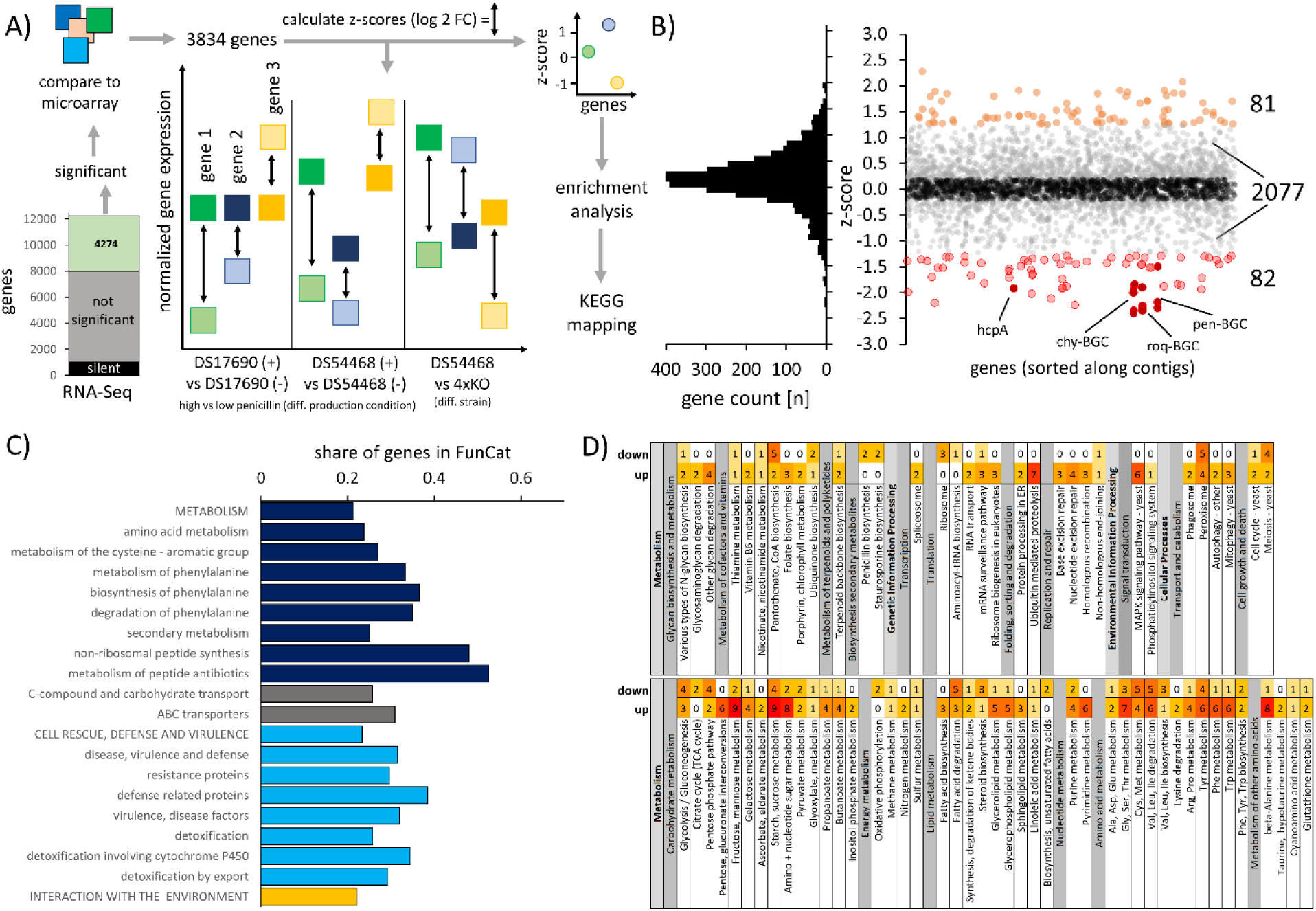
Analysis of transcriptome changes in 4xKO compared to DS54468 when growing in a glucose limited chemostat at a dilution rate of 0.05 h^-1^. A) Scheme for identification of genes that are differently expressed due to the absence of four BGCs. If the log2 FC of a gene in 4xKO and DS54468 is more different than observed log2 FCs from microarray experiments of strains grown in absence or presence of penicillin sidechain precursor, the z-score will become negative or positive, depending on the direction of the change. B) Distribution of z-scores for sufficiently covered genes and visualization of z-score over contigs, sorted from 1 to 49. Orange and red dots represent genes with a significantly different z-score (rate of false positives < 0.05, based on random sampling of normally distributed numbers). A clustering effect of negative z-score is seen for hcpA, chy-, roq- and pen-BGC which are highlighted. C) Enriched FunCat categories (p < 0.05, FDR corrected) derived from 2440 genes where |z| > 0.2. D) Overview of KEGG-pathways of *P. rubens* Wisconsin 54-1255 with up- and downregulated expression identified in this study.

Indeed, this gene set contained several genes that were involved in synthesis of the aromatic amino acids tryptophan, tyrosine and phenylalanine, which showed an increased abundance in the 4xKO strain. The entry reaction into the shikimate pathway is mediated by the 3-deoxy-7-phosphoheptulonate synthase (Pc18g02920, log2 FC 0.5) and subsequent enzymes showed also increased expression such as the anthranilate synthase (Pc13g12290, log2 FC 0.4). Further, the fate of aromatic amino acids differs, as the expression of the tryptophan synthase (Pc22g00910 log2 FC −1.4) is decreased in 4xKO, but further conversion of tryptophan to nicotinic acid is presumably enhanced because of increased expression of the kynureninases Pc22g20570 and Pc22g11870 (log2 FC 0.3 and 0.3, respectively) as well as the 3-hydroxyanthranilate 3,4-dioxygenase Pc20g09330 (log2 FC of 0.2). A decreased expression of tyrosinase (Pc22g18500, log2 FC −2.4) was noted which is presumably involved in formation of melanin. The second reaction in the catabolic route of tyrosine, the conversion of 4-hydroxyphenylpyruvate into homogentisate was upregulated with increased expression of the 4-hydroxyphenylpyruvate dioxygenase Pc22g07130 (log2 FC 0.4). The observed increase of phenylalanine levels seems to trigger an increased expression of Pc12g09020 (log2 FC 0.5), coding for a maleylacetoacetate isomerase, which is involved in phenylalanine degradation.

Also, an increased degradation of purines by adenosine deaminase (Pc06g00210, log2 FC 2.7) and guanine deaminase Pc16g11230 (log2 FC 0.6) might occur, while the adenine deaminase (Pc16g10530) showed decreased expression (log2 FC −0.5). Further downstream reactions in this pathway were also found to be upregulated, such as expression of IMP dehydrogenase (Pc13g07630, log2 FC 1.8), IMP and pyridine-specific 5’-nucleotidase (Pc12g13510, log2 FC 0.5) and xanthine dehydrogenase (Pc22g06330, log2 FC 0.6), while expression of the putative urate oxidase encoded by Pc22g20960 was decreased (log2 FC of −1.3). The adenosyl homocysteinase encoded by Pc16g05080 was lower expressed (log2 FC −0.3), putatively providing less adenosine. Also, expression of genes involved in pyrimidine biosynthesis was changed, namely the lower expression of the 4-amino-5-hydroxymethyl-2-methylpyrimidine phosphate synthase THI5 homolog encoded by Pc21g15700 (log2 FC −0.7) and an increased expression of the cytosine deaminase Pc16g10090 (log2 FC 0.5).

The biosynthesis of ribose 5-phosphate (R5P), the precursor required for purine biosynthesis and the pentose phosphate pathway was also found to be downregulated as the ribose 5-phosphate isomerase encoded by Pc22g21440 showed decreased expression (log2 FC −0.8). The link between the pentose phosphate pathway and glycolysis was also decreased as the Pc21g16950-encoded transaldolase was lower expressed (log2 FC −0.4), presumably resulting in an accumulation of D-arabitol, as the NADP+-dependent D-arabinitol dehydrogenase Pc16g08460 showed increased expression (log2 FC 1.2). Interestingly, it was found that the 6-phosphofructo-2-kinase Pc20g01550 (log2 FC 0.5) also showed increased expression. This is insofar interesting as the product, fructose-2,6-bisphosphate, strongly activates glycolysis through allosteric activation of phosphofructokinase 1. Further, we also found decreased expression of 2 pyruvate decarboxylase enzymes, encoded by Pc18g01490 (log2 FC – 0.7) and Pc13g09300 (log2 FC −0.6) which break down pyruvate into acetaldehyde and carbon dioxide, during anaerobic fermentation. Also, the carbonic anhydrase Pc22g06300 (log2 FC −0.6) was downregulated, which indicates that there might be less induction of these enzymes in the 4xKO strain.

Regarding the decrease in methionine and cysteine, this can perhaps be explained by decreased sulfur supply, as the phosphoadenosine phosphosulfate reductase Pc20g03220 was lower expressed (log2 FC - 0.5). This enzyme is required for fixation of inorganic sulfur via sulfite on 3’-phosphoadenosine-5’-phosphosulfate (PAPS). In addition, an increase of the cysteine dioxygenase expression (Pc21g04760, log2 FC 1.7), initiating the conversion of cysteine into taurine or degradation to sulfate was observed and might cause a decrease in methionine. Moreover, the cystathionine gamma-lyase (Pc21g05430, log2 Fc −0.8) breaking down cystathionine into cysteine, α-ketobutyrate and ammonia was downregulated. The lack of methionine might have been sensed by the cell as the L-methionine (R)-S-oxide reductase Pc20g05770 showed decreased expression (log2 FC −0.4). Since methionine and the related S-adenosylmethionine (SAM) act as cellular nutrient state sensors and have a significant impact upon regulation of autophagy^43,44^, the decrease in methionine levels also explains the upregulation of 11 autophagy-associated genes (log2 FC between 0.25 and 0.47, **Supplementary Information SI16**), amongst them the well-expressed (497.3 RPKM in DS54468) autophagy-related protein 8 (Atg8), encoded by Pc12g05370 (log2 FC 0.26). Atg8 is required for the formation of autophagosomal membranes during macroautophagy^45^ and is coupled to phosphatidylethanolamine (PE) by the cysteine protease Atg4^46^ (Pc20g08610, log2 FC 0.28). An increased expression of these genes is already observed when lowering the penicillin biosynthesis burden, as indicated by |z| < 0.2 (**Supplementary Information SI13)**. One exception is Pc12g10930 (z = −0.31, log2 FC of 0.47), encoding for the autophagy-related protein 13 (Atg13) that interacts with Atg1^47^ and whose phosphorylation status mediates interactions with other autophagy related proteins^45^. Here, an increased expression of *atg13* might suggest a further increase in autophagy as compared to strains still capable of producing penicillin. This could also result in an increased degradation of peroxisomes, which has been shown to be regulated by *atg1* in *P. rubens*^48^.

Taken together, these changes in gene expression are consistent with a reduced cellular demand for amino acids in the absence of the four highly expressed BGCs. It appears that the 4xKO cells redirect gene expression to avoid accumulation of aromatic amino acids. Also, since tryptophan and adenine are normally required for synthesis of NAD and NADP, having an increased pool under high-penicillin-production conditions^49^ is beneficial. However, these reservoirs trigger increased expression of degradation enzymes when demands are decreased as observed in the 4xKO strain. Overall, these changes suggest that the obtained strain will be suitable for the integration of heterologous gene clusters and expression thereof. To verify this, we next set out to overexpress endogenous and integrate heterologous secondary-metabolite related genes.

### *P. rubens* platform strain is suitable for multi-part donor DNA assembly and achieves increased yield of PKS-derived YWA1

To evaluate the secondary metabolite deficient strain as a platform, we first re-introduced the pen-BGC into the intergenic region (IGR) between Pc20g07090 and Pc20g08100 using *in vivo* homologous recombination using up to 8 DNA fragments with only 100 basepair overlap between fragments. We used two strategies for this approach, maintaining the native configuration of the cluster or replacing the promoters by the stronger p40s^50^ promoter (**Figure 5a**). While the first strategy led to a higher transformant number and successful integration, we observed a higher frequency of multiple gene copies being integrated when p40s was used, suggesting that recombination of very similar parts is less successful and prone to errors (**Figure 5b and Table 1**). The recombination was successful as evidenced by the concentration of penicillin V after five days of growth in SMP medium supplemented with phenoxyacetic acid (POA) (**Figure 5c**), even though the concentration of penicillin V was slightly lower for the recombined strains as compared to DS56830-penBGC and the reference strain for single-copy Pen-BGC production, DS54468. Investigation of the expression of all pen-BGC genes revealed high levels of expression in DS56830-penBGC and the strains expressing the pen-BGC genes from the p40s promoter (**Figure 5d**). The pen-BGC was similarly expressed in the 4xKO strain compared to the parental strain DS54468. Our results indicate that we can re-assemble a complete BGC from up to 8 DNA fragments with short homology successful, allowing efficient BGC pathway assembly with high functionality.

**Figure 4:**
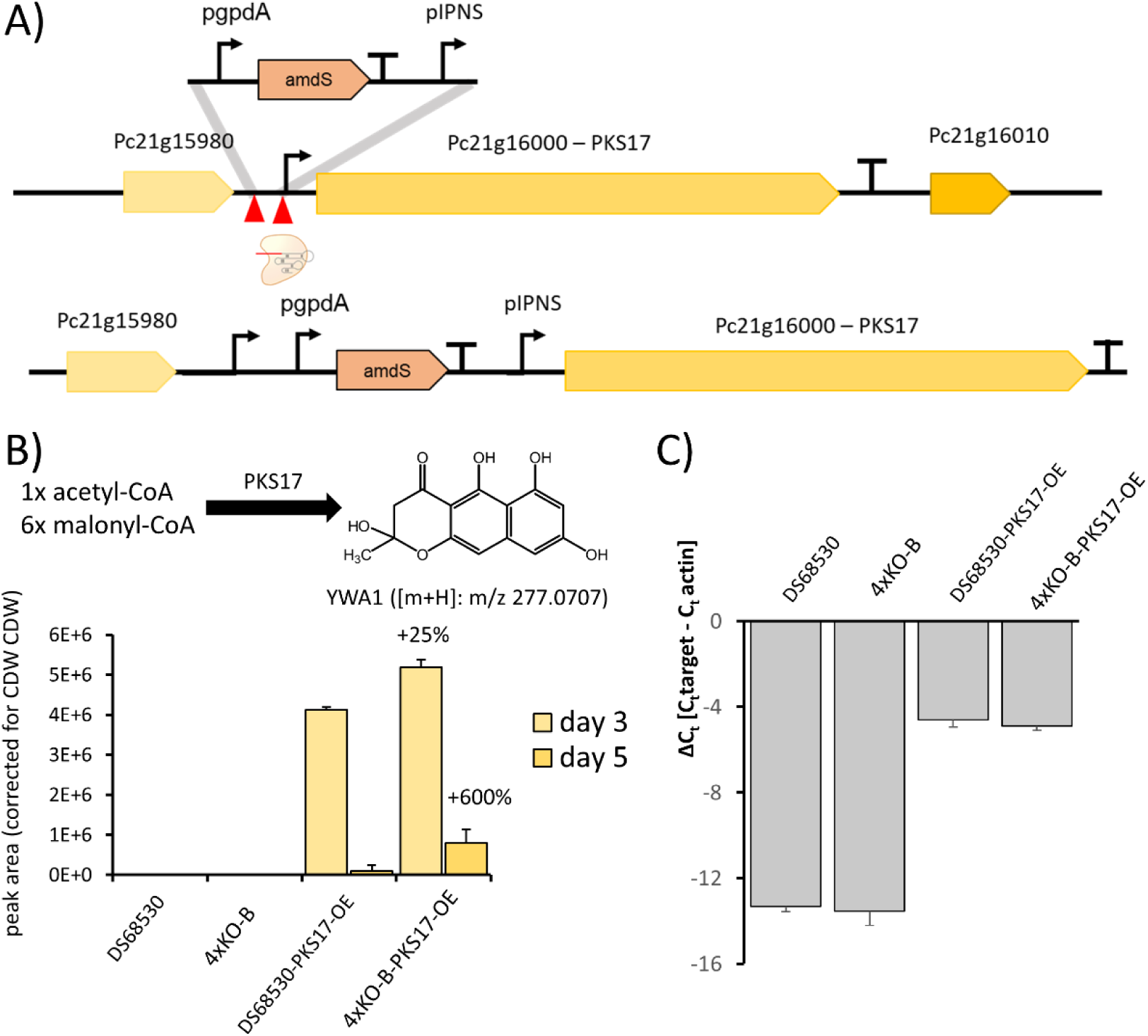
Overexpression of PKS17 in DS68530 and 4xKO-B. a) Schema showing the strategy used for integrating the IPNS promoter in front of Pc21g16000. b) The initial molecule produced by PKS17 is the naphthopyrone YWA1, which was quantified by LC-MS in fermentation broth of the indicated strains after 3 and 5 days. c) Expression of Pc21g16000 quantified by means of qPCR on day 3 of growth in SMP. The gene is not expressed in DS68530 and 4xKO-B and expression when replacing the promoter is unchanged between DS68530-PKS17-OE and 4KO-B-PKS17-OE, as seen by the difference in the similar Δct.

**Figure 5:**
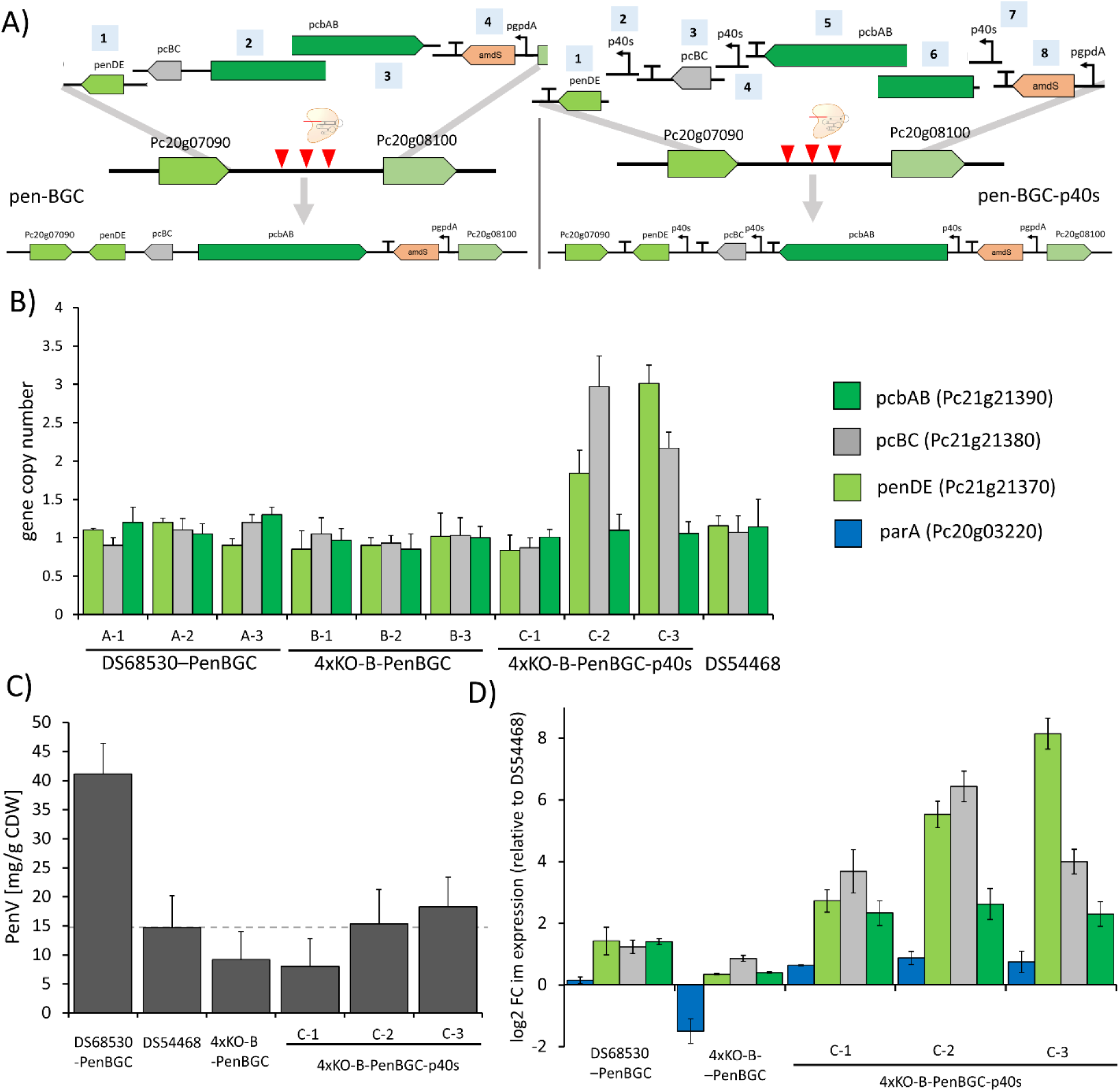
Integration of the Penicillin cluster into DS68530 and 4xKO-B. A) Scheme for recombination of parts obtained by PCR into the intergenic region of Pc20g07090 and Pc20g08100 using either the native promoters or p40s for expressing all genes of the pen-BGC. B) Copy number of integrated pen-BGC genes in the obtained strains. C) Penicillin V concentration after five days of growth in SMP + POA medium D) Changes in gene expression for pcbAB, pcBC, penDE and parA relative to the single-copy pen-BGC strain DS54468.

To further explore the performance of the platform strain, the polyketide synthase PKS17 (Pc16g1700) which is not expressed in *P. rubens* under submerged cultivation conditions was overexpressed by placing it under control of the pIPNS promoter (**Figure 4a**). Pc16g1700 encodes an iterative, non-reducing type I PKS, termed PKS17 or *pcAlb1*, producing the heptaketide naphthapyrone YWA1 by condensation of one acetyl-CoA and six malonyl-CoA moieties. YWA1 is the precursor for dihydroxynaphthalene (DHN)-melanin in several fungi^51^ and the BGC encoding the required enzymes is present in *P. rubens*^52^. Replacing the native promoter by pIPNS resulted in production of YWA1 in both DS68530-PKS17-OE and 4xKO-B-PKS17-OE, whereby the latter strain background increased the production of YWA1 by 25% on day 3 and by 600% on day 5 (**Figure 4b**). qPCR was used to measure expression of Pc16g1700 on day 5, and this showed no significantly different expression between DS68530-PKS17-OE and 4xKO-B-PKS17-OE (**Figure 4c**). Although YWA1 is further processed into insoluble pigments which were not quantified further, these results indicate that the supply of precursors for YWA1, malonyl-CoA and acetyl-CoA could be increased in the 4xKO background strain. Indeed, data from chemostat cultivations of 4xKO showed that Pc13g03920, encoding the *P. rubens* ortholog for the *S. cerevisiae* acetyl-CoA carboxylase (ACC) *Sce*ACC^53^ (UniProt Q00955.2) displayed a moderate increased expression (log2 FC of 0.36). ACC catalyzes the rate-limiting step in fatty-acid biosynthesis, the carboxylation of acetyl-CoA to malonyl-CoA, which is the limiting substrate for the biosynthesis of fatty acids via fatty-acid synthase^54^. Since the biosynthesis of YWA1 presents a drain for malonyl-CoA, an increased supply of malonyl-CoA via ACC could explain the increased level of YWA1. Our experiments show that the obtained 4xKO strain is suitable for producing increased amounts of the polyketide YWA1.

### Integration of heterologous Calbistrin gene cluster into the *P. rubens* platform strain results in production of decumbenones

To demonstrate that the obtained 4xKO platform strain is suitable for integration of heterologous BGCs, we integrated the calbistrin-BGC (cal-BGC) recently identified in *Penicillium decumbens*^55^. The products of the Cal-BGC are calbistrins and decumbenones, with the latter known to impair melanization of *Magnaporthe grisea*^56^, the cause of rice blast. Integration of the Cal-BGC into the genome of *P. rubens* 4xKO-B (**Table 1**) was achieved via *in vivo* homologous recombination (**Figure 6a and Supplementary Information SI17**). After liquid cultivation, we detected the linear moiety (dioic acid), decumbenone A, B and C but no Calbistrin A, C and versiol in both CYA and SMP medium samples of 4xKO-B-calBGC (**Figure 6b and Supplementary Information SI18**). Since our previous study could not rule out the possibility of a second PKS producing the linear moiety required for synthesis of calbistrins, we examined *P. rubens* for a possible upregulation of closely related PKSs with a potential homolog present in the calbistrin producers *A. versicolor, A. aculeatus, P. decumbens* and *C. tofieldiae*. The qPCR measurements did reveal a moderate increase in gene expression for Pc16g04890 (log2 FC of 1.3, **Supplementary Information SI19**), containing a C-methylation domain and an a enoyl-reductase domain, structurally resembling a highly-reducing PKS^57^ proposed to be necessary for synthesis of the linear moiety, however the confirmation of this hypothesis requires further experimental validation. Since we did not observe production of calbistrin A and C, this could suggest either a non-clustered broad specificity transesterase forms the ester bond of the calbistrins or hydrolysis of calbistrins is occurring very rapidly in *P. rubens*.

**Figure 6:**
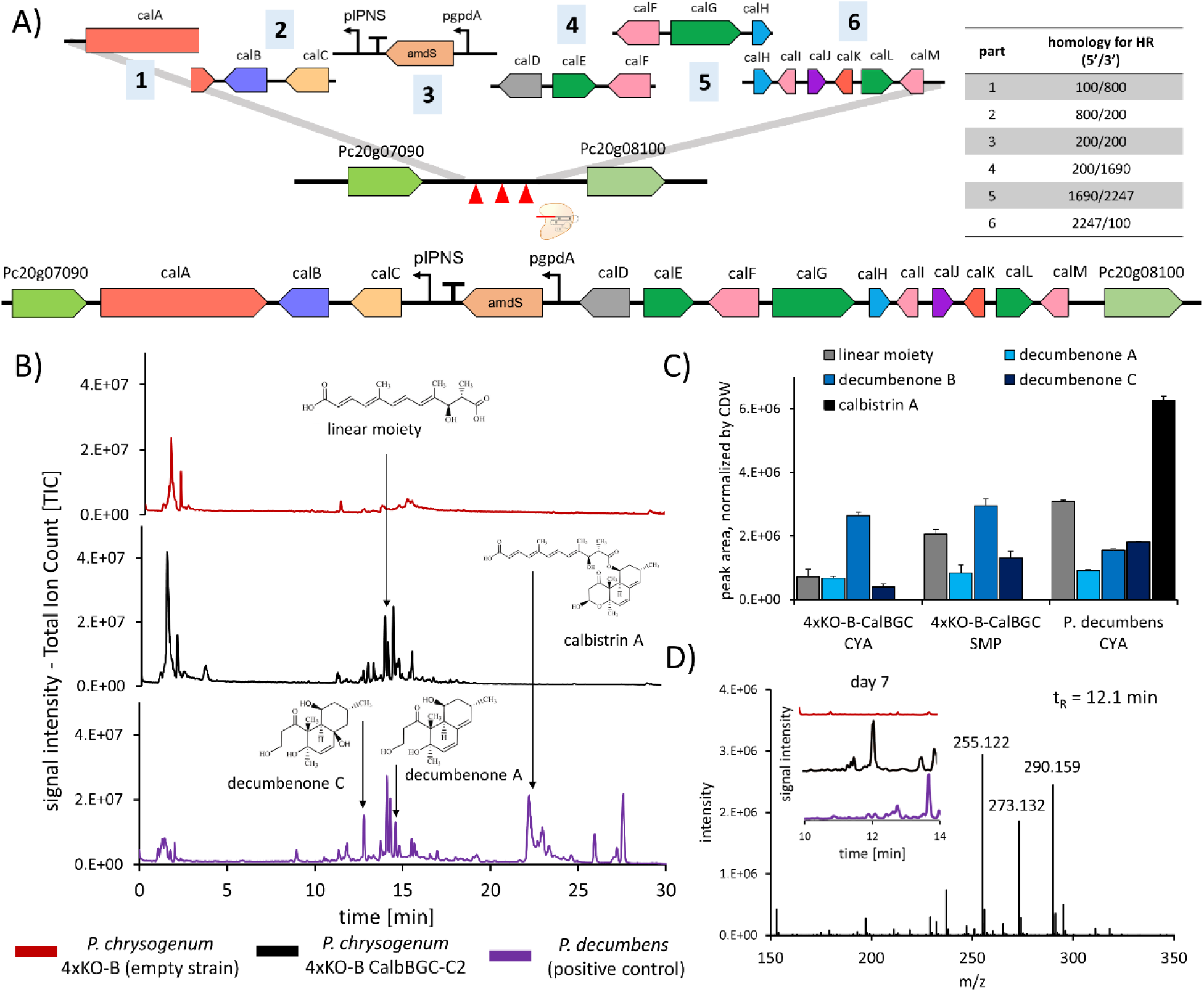
Integration of the calbistrin cluster from *P. decumbens* into 4xKO-B and verification of production. A) Scheme for recombination of six parts obtained by PCR into the intergenic region of Pc20g07090 and Pc20g08100. Obtained clones were verified by colony PCR (**Supplementary information 19**). B) Total-ion-chromatograms of samples taken five days after inoculation of CYA medium. Shown are 4xKO-B, a representative clone (4xKO-B-CalBGC-C2) and *P. decumbens*, serving as a positive control. Arrows indicate the retention times of the depicted molecules. C) Peak areas of calbistrin-related metabolites quantified in SMP medium and CYA medium taken five days after inoculation. Peak areas are depicted as mean of biological triplicates for 4xKO-B-CalBGC and biological duplicates for *P. decumbens*. No calbistrin A and C were detected in supernatant of 4xKO-B-CalBGC. An overview of retention times used m/z for quantification and obtained culture dry weight can be found in (**Supplementary Information 20 and 22**). D) Appearance of previously not observed peak in 4xKO-B CalBGC after 7 days of cultivation in CYA medium at a retention time of 12.1 min. Most abundant m/z in this peak were m/z 255.122, 273.132 and 290.159.

Expression of the Cal-BGC reduced biomass productionof P. rubens cultured in SMP medium and also abolished spore pigmentation (**Supplementary Information SI20**). However, for 4xKO-B-calBGC, the level of all decumbenones were increased in SMP medium at least one-fold when compared to CYA (**Figure 6c and Supplementary Information SI18**) with maximum levels observed on day 5 of growth. Besides reduced amounts of decumbenones, we also observed appearance of a significant peak on day 7 in 4xKO-B-calBGC, composed of 3 m/z: 255.122, 273.132 and 290.159 (**Figure 6d**), not observed in *P. decumbens*, suggesting degradation of decumbenones. Taken together, these observations indicate that the Cal-BGC contains all relevant genes for production of decumbenones and these can be successfully transferred to *P. rubens* for heterologous expression and high-level production.

## Conclusions

In this study, the consecutive deletion of well-expressed BGCs led to a secondary metabolite deficient strain of *P. rubens* that is suitable for integration and *in vivo* assembly of heterologous BGCs. By the use of *in vivo* homologous recombination employing multiple DNA fragments, a complete BGC can be reassembled while introducing at the same time promotors to enhance the expression. This methodology speeds up fungal synthetic biology leading to more freedom in the design-build-analyze-cycle. A major advantage of the platform strain is that novel heterologous compounds can be purified with reduced interference from endogenous secondary metabolites. We demonstrated this approach by heterologous expression of the calbistrin BGC from *Penicillium decumbens*, obtaining the melanization-inhibiting decumbenones as final products.

During the construction of the platform strain, intermediate strains were obtained with a different set of highly expressed NRPS genes. Metabolic profiling revealed an interesting interplay between the various NRPS enzymes. Because the substrate requirements of the enzymes PcbAB, ChyA, RoqA and HcpA show a certain degree of overlap, they likely compete for substrates and thus deletion of one BGC can result in higher levels of metabolites produced by the other BGCs. For instance, chrysogine biosynthesis seems to prevent accumulation of both roquefortine and fungisporine-related molecules by acting as a sink for anthranillic acid, the precursor for tryptophan biosynthesis. Since the chrysogine BGC is expressed under conditions required for penicillin biosynthesis, a deletion of the chrysogine BGC in a Pen-BGC strain might lead to a decreased penicillin production and shift the metabolic fingerprint to roquefortine-related molecules. The low expression of the remaining SM core genes in the 4xKO strain created here contributes to the clean secondary metabolite profile as indicated by the LC-MS data and adds to the applicability of this strain as a platform for secondary metabolism.

Our study also suggests that the demand for cysteine due to penicillin production naturally increases sulphate uptake via SutA/SutB and thus ensures higher methionine levels that decrease the autophagy response. The majority of amino acids did not display severe intracellular changes under the conditions utilized here, except methionine and nicotinic acid, the degradation product of tryptophan, suggesting that CSI of *P. rubens* towards an increased yield of ß-lactams did not result in a major impact on the cells ability to regulate amino acid metabolism by either reducing synthesis (methionine) or increasing degradation (tryptophan) of excess amino acids. Except for an extracellular increase of valine, other amino acid levels did not change drastically, hence the metabolism remains sufficiently flexible after CSI to respond to a decreased demand of certain amino acids. This will make the strain characterized here suitable for expression of both PKS- and NRPS-containing BGCs.

## Methods

### Data availability

High-throughput sequencing data have been deposited to the NCBI Sequence Read Archive database under accession PRJNA588889 (including SRA data SRR10428545, SRR10428546, SRR10428547, SRR10428548, SRR10428549, SRR10428550 and SRR10428551).

### Fungal Strains

All *P. rubens* strains utilized and created in this study can be found in (**Table 1**). Parental strains DS54468, DS68530 and DS47274 were kindly provided by DSM Sinochem B.V., now Centrient BV. *P. decumbens* strain IBT11843 was obtained from and is available at the IBT culture collection (Department of Biotechnology and Biomedicine, Technical University of Denmark).

### Nucleic Acid Techniques

If not indicated otherwise, PCRs were conducted using KAPA HiFi HotStart ReadyMix (Roche) and 0.25 µl of 100 µM primer stock solution at an annealing temperature of 66 °C regardless of calculated primer melting temperature and 30 seconds elongation per 1 kbp. PCR products were purified using the GenElute kit (Sigma Aldrich) and concentrations were checked using a NanoDrop ND1000 (Thermo Fisher Scientific).

For amplification of donor DNA parts, the origin of PCR templates is listed in **Supplementary information SI21**. For design of nucleic acid constructs and inspection of Sanger sequencing results, SnapGene (GSL Biotech) was used. For isolation of gDNA, the E.Z.N.A. HP Fungal DNA kit (Omega Biotek) was used to extract gDNA from 200 to 300 mg of wet or dried fungal biomass.

For qPCR analysis, mycelium for RNA extraction was collected by filtration, washed with ice-cold H_2_O. 100 to 200 mg wet biomass were mixed with 1 ml TRIzol reagent (Thermo Fisher Scientific), transferred into tubes containing glass beads and stored at −80 °C until RNA isolation. Mycelium was disrupted with a FastPrep FP120 system (Qbiogene) and total RNA was isolated using the Direct-zol RNA MiniPrep Kit (Zymo Research). For cDNA synthesis, 1500 ng total RNA were reverse transcribed using the Maxima H Minus cDNA Synthesis Master Mix (Life Technologies) in a volume of 20 µl. Samples were diluted with 80 µl MQ-H_2_O and 4 µl were used as input for qPCR in a final volume of 25 µl. As master mix for qPCR, the SensiMix SYBR Hi-ROX (Bioline Reagents) was used. All runs were performed on a MiniOpticon system (Bio-Rad). The following conditions were employed for amplification: 95 °C for 10 min, followed by 40 cycles of 95°C for 15 s, 60°C for 30 s and 72°C for 30 s, following an acquisition step. Raw c_t_ data were exported and analysis of relative gene expression was performed with the 2−ΔΔCT method^58^. The expression analysis was performed with two technical duplicates per biological sample. The γ-actin gene (Pc20g11630) was used as internal standard for data normalization. Primers for qPCR were designed using NCBI Primer BLAST^59^. All primers used for qPCR are listed in **Supplementary information SI21**. When appropriate, copy numbers of integrated donor DNA were analyzed using qPCR with extracted genomic DNA as an input according to Polli et al^60^.

RNA for RNA-seq was isolated from chemostat-cultured mycelium which was obtained by sampling 10 ml broth, vacuum filtration and brief washing with water before freezing in liquid nitrogen and storing at −80°C until RNA extraction. For RNA extraction, mycelium was ground in liquid nitrogen followed by TRIzol-chloroform and isopropanol treatment to precipitate total RNA. A further purification step was done using the Nucleospin RNA clean-up Kit (Machery-Nagel), following storage of samples at −80 °C until shipping for RNA-seq analysis on dry ice. From each sample, 10 µg were submitted for generation of 2×10^6^ 50 bp single-end reads (50 bp) on a BGISEQ-500 (BGI).

### Fungal Techniques

Protoplasts of *P. rubens* strains were obtained 48 hours post spore seeding in YGG medium and transformed using the methods and media described previously^25,37^. The amount of total DNA transformed did not exceed 10 µg in a maximum volume of 50 µl. A list of all conducted transformations during this study is given in **Table 1**.

Selection was carried out by utilizing transformant recovery plates containing either 0.1% acetamide (Sigma Aldrich) as only nitrogen source or 50 μg/ml phleomycin (Invivogen) or 1.2 μg/ml terbinafine hydrochloride (Sigma Aldrich) combined with 40 mM sodium nitrate as nitrogen source. After transformants recovered, colonies were counted and analyzed by colony PCR using the Phire Plant Direct PCR Kit (Thermo Fisher Scientific) to confirm integration of donor DNA elements at the desired genomic locus.

When appropriate, PCR products were purified using ExoSAP-IT PCR Product Cleanup Reagent (Thermo Fisher Scientific) and sent for sequencing (Macrogen) with suitable primers. Correct clones were purified by two to three cycles of sporulation and subsequent spore plating on selective media. For sporulation, fungal cultures where maintained on R-Agar^61^ for 7 to 10 days.

For long-term storage of strains, rice batches were prepared, lyophilized and stored at room temperature when used for inoculation of shake flask cultures. For long-term cryocultures, rice grains with attached spores were stored in 20% glycerol solution at −80°C.

All shake flask cultivations were performed in 100 ml flasks shaken at 200 rpm and 25°C in a shaking incubator. For precultures, 25 ml YGG medium^61^ was inoculated with lyophilized rice grains (between 0.2×10^6^ to 2×10^6^ spores/ grain, using 1 grain per ml inoculum) and incubated for 48 h. Of that preculture 4 ml (without rice grains) was used to inoculate either 26 ml SMP^61^ or CYA^62^ medium. When appropriate, SMP was supplemented with 2.5 g/L of phenoxyacetic acid (POA) to stimulate production of Penicillin V. Samples were harvested by vacuum filtration over cellulose filters (Sartorius) at indicated times for biomass determination and RNA isolation. The filtrated broth was further clarified using 0.2 μm polytetrafluoroethylene (PTFE) syringe filter (VWR) and stored at −80°C until further analysis by HPLC analysis and LC-MS.

### Bioreactor Cultivations

Cultivation was done in 7.5 L BioFlo310 bioreactors (New Brunswick Scientific). Glucose-limited chemostat cultivation was initiated by inoculation of 5 L ammonium based minimal medium (MM) pH 6.5 as described by Douma et al.^63^. No sidechain precursors for PenV or PenG production were added. Inoculations were performed with a conidial suspension to give 10^9^ conidia/L cultivation medium. Germination was facilitated by addition of 0.003% (w/w) yeast extract. Temperature of 25°C and pH 6.5 were kept constant, the latter by controlled addition of 2 M NaOH or 1 M HCl, respectively. After an initial germination of 15 hours, the stirrer speed was increased from 500 rpm to 750 rpm and aeration with air was changed from head space to sparger gassing (1L/min). Subsequently, polypropylene glycol 2000 (PPG) at a final concentration of 0.01% (v/v) was added as an antifoaming agent. Continuous cultivation was started in the late exponential growth phase, when 100 ml of 2 M NaOH had been added to the batch culture (biomass concentration of about 6 g dry weight per kg of culture)^64^. MM containing 0.01% (v/v) PPG was fed to the culture from two interconnected 20-liter reservoirs. The flow rate was 0.25 L/h, which corresponded to a dilution rate (D) of 0.05 h-1. The weight was kept constant at 5 kg using a port at the bottom of the reactor. The outflow was regulated by a magnetic valve operating at a band width of 10 g. Steady-states were defined by constant alkali addition rate, produced CO2, consumed O2 and biomass concentration after more than four residence times (> 80 h). Samples were taken regularly and frozen quickly in liquid nitrogen. Mycelium harvested during steady-state conditions was used for RNA-seq analyses. Additional quadruplicate samples were taken directly into −20°C 40% methanol (20 mL) to determine intracellular amino acids. The content of CO_2_ and O_2_ in the exhaust gas was analyzed using an Ex-2000 Gas Analyser (New Brunswick Scientific). The pH was measured with an autoclavable glass electrode (Mettler Toledo), and the dissolved oxygen tension was measured with an lnPro-6860i O2 sensor (Mettler Toledo).

### Amino acid quantification

Mycelium for amino acid analysis was harvested during steady-state according to the method described by de Jonge et al^65^. In brief, samples of 10 ml or less were quenched directly into −20°C 40% methanol (20 ml), weighted and filtered using a vacuum pump followed by a single washing 1x with the same volume of ice-cold 40% methanol before freezing in liquid nitrogen and storage at −80°C until extraction. For extraction, filter papers with frozen sample were directly placed in 50 ml falcon tubes containing 20 ml 73°C hot 75% ethanol, shaken vigorously, boiled for 3 min at 95°C, chilled on ice for 5 min, centrifuged for 5 min at 4000 x g^-1^ and filtered over a 0.2 µm cellulose acetate filter (VWR). 1 ml aliquots were concentrated in a speed-vac (Eppendorf) for 45 min at 30°C, centrifuged for 10 min at 10.000 x g^-1^. Supernatant was stored at −80°C if not used immediately for LC-MS analysis. All extractions were performed in quadruplicate per bioreactor run and analyzed in technical duplicate on LC-MS. Amino acid retention times were verified by a standard mixture (AAS18 Analytical standard; Sigma Aldrich) or dilutions of pure amino acids in 10 mM HCl (for Asn, Gln, Trp). Peak areas were corrected for extracted biomass and concentrations were calculated using a calibration curve.

### LC-MS analysis

For analysis of broth, samples were centrifuged for 10 min at 14,000 x g^-1^ and supernatant was directly used for analysis or aliquoted and frozen at −80°C. Separation was conducted on an Accella1250 UPLC system coupled to an Orbitrap Exactive spectrometer (Thermo Fisher Scientific, The Netherlands) with a scan range of m/z 100 – 1600 Da. A sample of 5 μL was injected onto a Shim-pack XR-ODS C18 column (3.0 × 75 mm, 2.2 μm ID) (Shimadzu, Japan) kept at 40 °C and operated at a flow rate of 300 μL/min. Separation of compounds was achieved with the following solvents: A: 100% MQ-H2O, B: 100% Acetonitrile, and C: 2% formic in MQ-H2O being constantly added at 5% to protonate molecules. After sample injection, the column was run for 5 min with isocratic flow at 5 % B, following a linear gradient for 25 min reaching 95% B, remaining constant at 95% B for 5 min and equilibrating the column with initial conditions of 5% B for 5 min before injection of the next sample. Each sample was analyzed in technical duplicate. Retention times for N-acetylglucosamine, nicotinic acid and amino acids were verified with available standards (purity 99%, Sigma-Aldrich). Peak areas were extracted with Thermo Xcalibur (Version 2.2 SP1) software allowing a Δm/z of 5 ppm to the exact mass of compound m/z. Chromatogram data in Thermo-RAW format were imported into XCMS^38,66^ and processed with default settings to identify enriched and depleted m/z.

### HPLC-Analysis

The extracellular concentration of Penicillin V was determined according to the method described by Harris et al^67^ using a Nextera HPLC system with a Shim-pack XR-ODS 2.2 RP column (Shimadzu). Isocratic separation was used for 1 µl of 0.2 µm filtered fermentation broth at a flow rate of 0.5 mL/min with running buffer (acetonitrile 245 g/L, 640 mg/L, KH_2_PO_4_ and 340 mg/L phosphoric acid in Milli-Q). Peaks were detected at a wavelength of 254 nm according to the retention time of standards between 11.5 and 12 minutes. Production levels were corrected for growth differences by dry weight determination. Samples were analyzed in technical duplicate.

### Transcriptome Data Processing

Mapping of raw reads was performed with DNAStar Lasergene 14 Suite using the *P. rubens* assembly GCA_000226395.1 as reference for alignment. For each biological replicate, the RPKM (Reads Per Kilobase Million) were calculated individually and replicates were averaged as the number of mapped reads was similar **(Supplementary Information SI22)**. P-values were calculated using a 2-sided students t-test assuming unequal variance for each gene. Genes with an RPKM below 0.3 were considered silent or potential noise of gDNA contamination. Genes that were silent (below 0.3 RPKM) in one strain but expressed above 0.3 RPKM in the other were assigned a log2 FC of 7/-7 respectively. PaintOmics3^68^ was used to generate overviews for pathways displaying changed expression patterns and for hypothesis generation of biologically relevant patterns. Statistically relevant genes were processed with FungiFun2^69^ to identify enriched functional annotation scheme (FunCat^70^) categories using FDR-correction and indirectly annotated top categories. For comparison of expression changes in high versus low penicillin production conditions across different strains, z-scores were calculated. Z-scores represent the distance between the raw score (log2 FC derived from RNA-seq data generated in this study) and the population mean (mean log2 FC derived from multiple high versus low penicillin production conditions) in units of standard deviation. Z-scores are negative for raw scores below the mean and positive when above. The log2 FC expression changes of significantly different expressed genes from this study (|log 2 FC| > 0.25) were extracted from published microarray data (**Supplementary Information SI1**, only genes covered by at least 50% of the input microarray data sets were considered). Assuming an error rate of 5% and a normal distribution of log2 FC values, the confidence interval for a significantly different expression between the conditions was calculated to be [-1.25; 1.25]. The calculated z-scores of genes were plotted over contigs of *P. rubens* aligned from 1 to 49 to identify possible clustering effects due to co-expression at the same genomic locus.

### Off-target and SNP-analysis

Possible off-target sites were identified using Cas-OFFinder^71^ and are listed in **Supplementary Information SI23**. OFF-target sites without a bulge and less than 5 mismatches were verified by PCR amplification of the locus followed by sanger sequencing (Macrogen). Additionally, isolated genomic DNA was sent to BGI Europe (Copenhagen, Denmark) for genomic library preparation and 100 bp paired-end sequencing. From each strain, above 10 million reads were generated and > 98% were successfully mapped to the *P. rubens* Wisconsin 54-1255 genome using Breseq^72^ to detect mutations.

### Phylogenetic Tree Construction for shared PKS in Calbistrin Producers

All proteins containing a PKS_KS superfamily domain were retrieved from NCBI via BLAST search for *P. rubens* Wisconsin 54-1255, *A. versiciolor* CBS 583.65, *A. aculeatus* ATCC 16872, *Colletotrichum tofieldiae* 0861 and *P. decumbens* IBT11843, yielding 137 proteins. Full-length-protein sequences were aligned using MUSCLE with default settings in MEGA X^73^ and evolutionary history was inferred using 500 bootstraps and a Jones-Taylor-Thornton (JTT) model for amino acid substitutions assuming uniform rates of mutations among all sites. A total of 67 amino acid positions were used in the final dataset for tree construction. The obtained tree was inspected for branches containing proteins from all input species and expression of candidate genes was analyzed by qPCR in *P. rubens* 4xKO-B strains expressing the Cal-BGC grown for 5 days in SMP medium.

## AUTHOR INFORMATION

### Author Contributions

RALB and AJMD designed the experiments and conceived the study.

CP, AV and FP performed gene deletions and shake cultivations of strains.

TS and CP performed bioreactor cultivations and sampling. CP and SJ extracted samples and performed analytical experiments.

CP and FP analyzed mass spectrometry data.

CP analyzed transcriptome data.

MdV performed integration of penicillin and calbistrin BGCs and performed shake-flask cultivations. LM performed shake flask cultivations, HPLC analysis and SNP data.

AV performed qPCR experiments and analyzed data.

CP and TS wrote the manuscript with the improvements from all authors. All authors read and approved the final manuscript.

## Acknowledgments

The research leading to these results was financially supported by the BE-Basic Foundation (Biotechnology based Ecologically Balanced Sustainable Industrial Consortium), DSM Sinochem Pharmaceuticals Netherlands B. V. and the European Commission’s Marie Curie Initial Training Network Quantfung (FP7-People-2013-ITN, grant no. 607332). The authors would like to thank Oleksandr Salo for technical assistance with LC-MS. Hilde E. Havinga is acknowledge for assistance with shake-flask cultivations and analysis of samples with HPLC. Sietske Grijseels and Rasmus N. Frandsen are acknowledged for valuable discussion about the calbistrin cluster. Jeroen G. Nijland is acknowledged for support with BreSeq and genome sequencing.

## Competing interests

All authors declare no competing financial interests.

