## Supplementary Data for "A *Penicillium rubens* platform strain for secondary metabolite production"

[3] Van Hall Larenstein University of Applied Sciences, Leeuwaarden, The Netherlands

A)

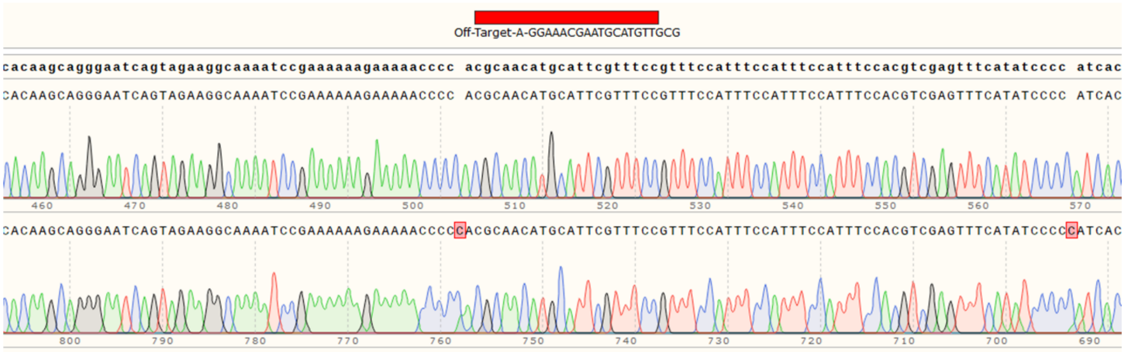
Off-Target-A-GGAAACGAATGCATGTTGCG

B)

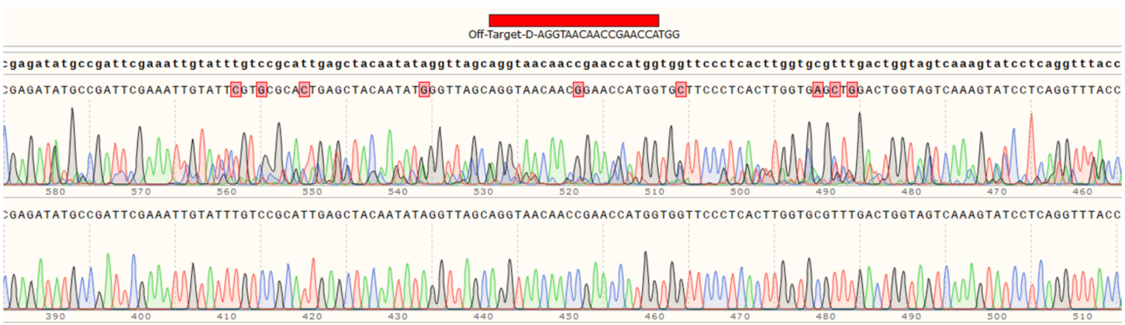
Off-Target-D-AGGTAACAACCGAACCATGG

C)

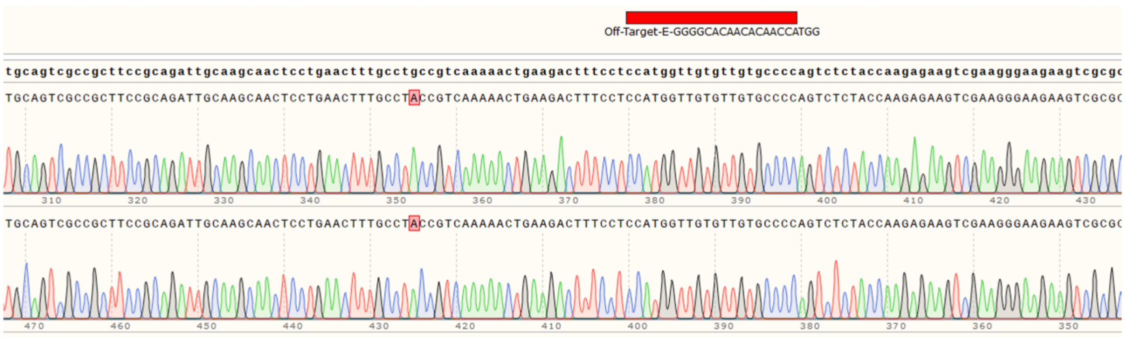
Off-Target-E-GGGGCACAACACAACCATGG

D)

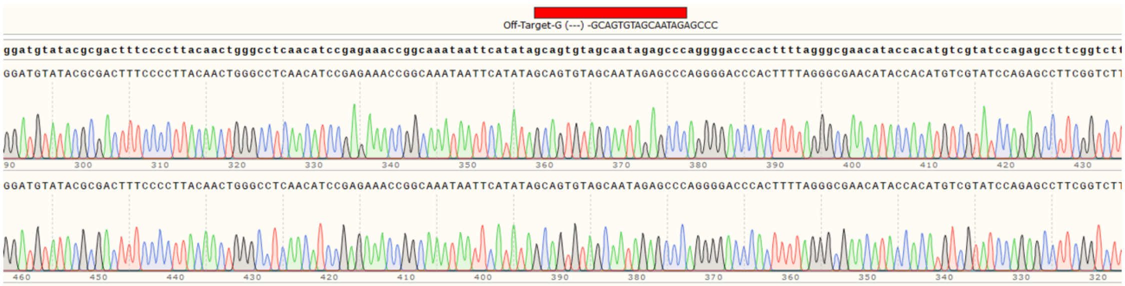
Off-Target-G (---) -GCAGTGTAGCAATAGAGCCC

E)

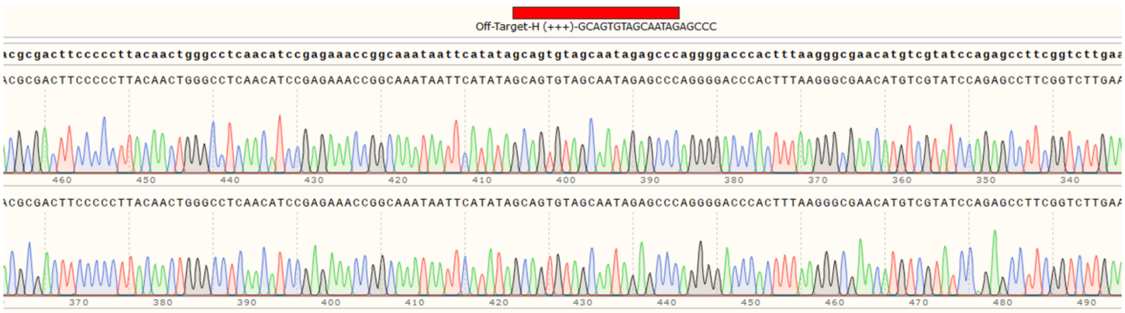
Off-Target-H (+++)-GCAGTGTAGCAATAGAGCCC

**Supplementary information SI4** – Results of Sanger sequencing of PCR products obtained from gDNA of 4xKO to verify absence of point mutations at off-target sites. A) Off-target A in terminator of Pc13g09530; B) Off-target D located in intron of Pc20g03630; C) Off-target E located in promoter of Pc21g03670; D) Off-target G in intron of Pc21g00490; E) Off-target H located in promoter of Pc24g01430. All sequencing results did not display any unexpected mutational event.

| group | compound | formula | m/z ([m+H] +/- 5 ppm ) | TR [min] | day | 2 | 3 | 4 | 5 | 6 | 7 |
|---|---|---|---|---|---|---|---|---|---|---|---|
| Penicillin | LLD-ACV | $C_{14}H_{25}N_3O_6S$ | 364.15 | 10.0 | | 4.0E+05 | 8.1E+05 | 3.8E+05 | 1.4E+05 | 2.6E+05 | 1.3E+05 |
| | IPN | $C_{14}H_{21}N_3O_6S$ | 360.12 | 10.0 | | 5.1E+03 | 7.9E+03 | 8.0E+03 | 1.0E+04 | 1.3E+04 | 2.2E+04 |
| | 6-APA | $C_8H_{21}N_2O_3S$ | 217.06 | 1.8 | | 8.1E+05 | 3.4E+06 | 5.9E+06 | 1.1E+07 | 1.1E+07 | 2.2E+07 |
| Chrysogine-associated | Chrysogine | $C_{17}H_{17}N_5O_2$ | 191.09 | 12.0 | Peak area in DS27472 (corrected for CDW) | 2.5E+07 | 2.4E+07 | 2.2E+07 | 2.4E+07 | 2.3E+07 | 2.1E+07 |
| | Chrysogine 2 | $C_{12}H_{15}N_3O_3$ | 250.12 | 11.2 | | 1.1E+06 | 2.3E+06 | 3.3E+06 | 3.6E+06 | 3.4E+06 | 3.6E+06 |
| | chrysogine 3 | $C_{10}H_{10}N_2O_3$ | 207.08 | 9.7 | | 4.6E+03 | 1.3E+05 | 4.4E+05 | 1.0E+06 | 2.1E+06 | 2.6E+06 |
| | Chrysogine 4 | $C_{13}H_{15}N_3O_5$ | 294.11 | 11.0 | | 9.8E+05 | 4.5E+06 | 1.0E+07 | 1.3E+07 | 1.4E+07 | 1.6E+07 |
| | Chrysogine 5 | $C_{15}H_{20}N_4O_5$ | 337.15 | 7.7 | | 1.2E+05 | 7.4E+05 | 1.3E+06 | 9.7E+05 | 1.2E+06 | 1.1E+06 |
| | Chrysogine 6 | $C_{15}H_{18}N3O_6$ | 336.12 | 9.0 | | 4.1E+02 | 5.3E+02 | 5.2E+02 | 7.2E+02 | 7.7E+02 | 7.1E+03 |
| | Chrysogine 7 | $C_{13}H_{12}N_2O_5$ | 277.08 | 10.0 | | 2.5E+05 | 1.2E+06 | 2.6E+06 | 3.2E+06 | 3.8E+06 | 4.4E+06 |
| | Chrysogine 8 | $C_{13}H_{14}N_2O_6$ | 295.09 | 11.4 | | 1.3E+05 | 6.3E+05 | 1.4E+06 | 1.8E+06 | 1.9E+06 | 2.1E+06 |
| | Chrysogine 9 | $C_{20}H_{20}N_4O_6$ | 413.14 | 15.0 | | 4.0E+04 | 3.2E+04 | 3.2E+04 | 3.1E+00 | 3.1E+04 | 2.9E+04 |
| | chrysogine 10 | $C_{20}H_{20}N_4O_6$ | 413.14 | 15.5 | | 2.4E+03 | 6.1E+04 | 6.2E+04 | 3.7E+04 | 4.8E+04 | 3.8E+04 |
| | Chrysogine 12 | $C_{13}H_{13}N_3O_4$ | 276.10 | 12.3 | | 3.3E+03 | 1.6E+05 | 2.1E+05 | 3.8E+05 | 5.7E+05 | 6.0E+05 |
| | Chrysogine 13 | $C_{15}H_{19}N_3O_6$ | 338.13 | 13.0 | | 1.3E+05 | 6.1E+05 | 1.0E+06 | 8.8E+05 | 7.4E+05 | 6.7E+05 |
| Roquefortine-associated | HTD | $C_{17}H_{17}N_5O_2$ | 324.145 | 4.5 | Peak area in DS27472 (corrected for CDW) | 7.8E+02 | 2.7E+05 | 4.0E+05 | 1.3E+06 | 2.6E+06 | 4.4E+06 |
| | DHTD | $C_{17}H_{15}N_5O_2$ | 322.129 | 5.7 | | nd | 2.1E+04 | 1.8E+04 | 3.5E+03 | 8.3E+04 | 1.4E+06 |
| | Roquefortine D | $C_{22}H_{25}N_5O_2$ | 392.208 | 16.3 | | 5.0E+04 | 1.1E+05 | 2.1E+05 | 5.0E+05 | 1.3E+06 | 1.4E+06 |
| | Roquefortine C | $C_{22}H_{23}N_5O_2$ | 390.193 | 21.0 | | nd | nd | 1.0E+02 | 2.1E+02 | 6.4E+02 | 3.9E+03 |
| | Glandicoline A | $C_{22}H_{21}N_5O_3$ | 404.172 | 16.4 | | nd | 3.2E+04 | 1.7E+04 | 1.9E+04 | 8.6E+04 | 4.3E+05 |
| | Glandicoline B | $C_{22}H_{21}N_5O_4$ | 420.167 | 17.0 | | nd | nd | nd | nd | nd | nd |
| | Meleagrin | $C_{23}H_{23}N_5O_4$ | 434.182 | 17.0 | | nd | 6.6E+04 | 2.5E+05 | 8.7E+05 | 5.1E+06 | 2.3E+07 |
| | Neoxaline | $C_{23}H_{25}N_5O_4$ | 436.198 | 17.0 | | nd | nd | nd | 3.2E+04 | 1.1E+05 | 4.0E+05 |
| | Roquefortine F | $C_{23}H_{17}N_5O_2$ | 420.203 | 19.0 | | nd | nd | nd | 3.5E+03 | 7.6E+03 | 3.0E+04 |
| | Roquefortine M | $C_{22}H_{23}N_5O_4$ | 422.181 | 16.0 | | nd | nd | nd | 1.0E+03 | 6.2E+03 | 1.9E+04 |
| | Roquefortine N | $C_{22}H_{25}N_5O_5$ | 440.192 | 11.80 | | nd | nd | nd | 9.9E+02 | 1.2E+03 | 2.3E+03 |
| Fungisporin-associated | cyclo-(YFVV) | $C_{28}H_{36}N_4O_5$ | 509.276 | 26.6 | Peak area in DS27472 (corrected for CDW) | 2.6E+04 | 1.5E+05 | 4.6E+04 | 1.5E+05 | 9.0E+04 | 1.9E+05 |
| | cyclo-(YWVV) | $C_{30}H_{37}N_5O_5$ | 548.287 | 26.6 | | nd | nd | nd | nd | nd | nd |
| | cyclo-(FFVV) - Fungisporin | $C_{28}H_{36}N_4O_4$ | 493.281 | 27.3 | | nd | nd | nd | nd | 2.9E+03 | 4.2E+03 |
| | cyclo-(YFIV) | $C_{29}H_{38}N_4O_5$ | 523.291 | 26.9 | | nd | nd | nd | nd | nd | nd |
| | cyclo-(FWVV) | $C_{30}H_{37}N_5O_4$ | 532.292 | 27.2 | | 3.0E+02 | 4.3E+03 | 4.3E+03 | 4.1E+03 | 7.3E+03 | 3.2E+04 |
| | cyclo-(FFVI) | $C_{29}H_{38}N_4O_4$ | 507.296 | 27.7 | | nd | nd | nd | nd | nd | nd |
| | cyclo-(YWVI) | $C_{31}H_{39}N_5O_5$ | 562.302 | 26.86 | | nd | nd | nd | nd | 1.1E+04 | 2.2E+04 |
| | YFVV, VYFV, FVVY | $C_{28}H_{38}N_4O_6$ | 527.286 | 20.1 | | nd | 1.3E+04 | 4.4E+04 | 8.0E+04 | 2.2E+05 | 9.4E+05 |
| | YWVV,VYWV, WVVY | $C_{30}H_{39}N_5O_6$ | 566.297 | 20.4 | | nd | 4.6E+03 | 2.3E+04 | 3.9E+04 | 7.5E+04 | 2.6E+05 |
| | FVIY, IYFV, FIVY, VYFI | $C_{29}H_{40}N_4O_6$ | 541.302 | 21.4 | | nd | nd | 4.1E+03 | 5.4E+03 | 9.7E+03 | 6.0E+04 |
| | VFWV, FWVV, WVVF | $C_{30}H_{39}N_5O_5$ | 550.302 | 23.6 | | 1.0E+04 | 1.6E+04 | 2.1E+04 | 7.3E+04 | 7.2E+04 | 2.3E+05 |
| | FVVF, VFFV, FFVV | $C_{28}H_{38}N_4O_5$ | 525.308 | 23.3 | | 1.5E+04 | 3.0E+04 | 6.2E+04 | 1.1E+05 | 2.1E+05 | 7.8E+05 |
| | FVIF, FIVF | $C_{29}H_{40}N_4O_5$ | 525.307 | 24.7 | | nd | nd | 6.4E+02 | 2.4E+03 | 5.2E+03 | 2.6E+04 |

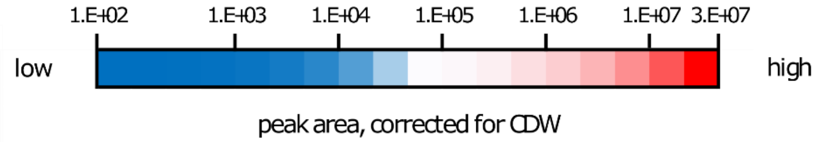


**Supplementary Information SI5:** Average peak areas normalized for cell dry weight (CDW) of molecules associated with penicillin, chrysogine, roquefortine and fungisporin biosynthesis in fermentation broth of parental strain DS27472 during cultivation in shake flasks using SMP medium. Samples were taken in 24 hour intervals starting on day 2 (n= 3 independent shake flasks). nd = not detected, retention times (TR) are indicated

| | | strain | DS68530 (ΔPen) | | | | | | DS68530 (ΔPen, ΔChy) | | | | | | DS68530 (ΔPen, ΔChy, ΔRoq) | | | | | | DS68530 (ΔPen, ΔChy, ΔhcpA) | | | | | |
|---|---|---|---|---|---|---|---|---|---|---|---|---|---|---|---|---|---|---|---|---|---|---|---|---|---|---|
| | compound | day | 2 | 3 | 4 | 5 | 6 | 7 | 2 | 3 | 4 | 5 | 6 | 7 | 2 | 3 | 4 | 5 | 6 | 7 | 2 | 3 | 4 | 5 | 6 | 7 |
| Chrysogine-associated | Chrysogine | change in peak are, relative to DS27472 | 2.7 | 1.5 | 1.4 | 1.5 | 1.1 | 1.2 | | | | | | | | | | | | | | | | | | |
| | Chrysogine 2 | | 1.2 | 1.2 | 1.1 | 1.2 | 1.3 | 1.2 | | | | | | | | | | | | | | | | | | |
| | chrysogine 3 | | 1.0 | 1.3 | 1.4 | 1.6 | 1.4 | 1.3 | | | | | | | | | | | | | | | | | | |
| | Chrysogine 4 | | 1.2 | 1.3 | 1.1 | 1.1 | 1.1 | 1.1 | | | | | | | | | | | | | | | | | | |
| | Chrysogine 5 | | 2.2 | 1.6 | 1.1 | 1.2 | 0.6 | 0.3 | | | | | | | | | | | | | | | | | | |
| | Chrysogine 6 | | 1.7 | 3.0 | 2.4 | 15.6 | 23.9 | 11.5 | | | | | | | | | | | | | | | | | | |
| | Chrysogine 7 | | 1.3 | 1.4 | 1.1 | 1.1 | 1.1 | 1.1 | | | | | | | | | | | | | | | | | | |
| | Chrysogine 8 | | 1.2 | 1.3 | 1.0 | 1.1 | 1.2 | 1.1 | | | | | | | | | | | | | | | | | | |
| | Chrysogine 9 | | 3.6 | 3.1 | 2.9 | 2.7 | 1.6 | 2.2 | | | | | | | | | | | | | | | | | | |
| | chrysogine 10 | | 29.0 | 4.1 | 3.8 | 3.5 | 3.0 | 2.8 | | | | | | | | | | | | | | | | | | |
| | Chrysogine 12 | | 4.2 | 1.3 | 0.9 | 1.1 | 0.8 | 1.0 | | | | | | | | | | | | | | | | | | |
| | Chrysogine 13 | | 1.5 | 1.1 | 0.8 | 0.6 | 0.3 | 0.4 | | | | | | | | | | | | | | | | | | |
| Roquefortine-associated | HTD | change in peak are, relative to DS27472 | 0.7 | 0.5 | 0.4 | 0.3 | 0.4 | 0.5 | 4.2 | 3.7 | 3.3 | 4.0 | 3.9 | 3.6 | | | | | | | 1.5 | 1.4 | 2.1 | 2.5 | 3.2 | 1.4 |
| | DHTD | | n/a | n/a | n/a | n/a | 0.2 | 0.3 | n/a | n/a | 0.0 | 0.1 | 0.2 | 0.5 | | | | | | | n/a | n/a | 0.1 | 0.0 | 0.0 | 0.0 |
| | Roquefortine D | | n/a | 0.3 | 0.2 | 0.3 | 0.1 | 0.2 | 4.3 | 4.5 | 4.7 | 7.6 | 10.4 | 15.3 | | | | | | | 1.9 | 2.1 | 2.0 | 3.2 | 2.4 | 1.7 |
| | Roquefortine C | | n/a | n/a | 5.7 | 3.6 | 1.4 | 1.3 | n/a | n/a | 4.9 | 4.8 | 2.5 | 2.6 | | | | | | | n/a | n/a | n/a | 9.4 | 6.0 | 4.7 |
| | Glandicoline A | | n/a | 0.0 | 0.1 | 0.1 | 0.1 | 0.5 | n/a | 0.1 | 0.5 | 12.6 | 4.5 | 1.5 | | | | | | | n/a | 0.2 | 0.4 | 14.5 | 7.7 | 1.6 |
| | Glandicoline B | | n/a | n/a | n/a | n/a | n/a | n/a | n/a | n/a | n/a | n/a | n/a | n/a | | | | | | | n/a | n/a | n/a | n/a | n/a | n/a |
| | Meleagrin | | n/a | 0.2 | 0.2 | 0.1 | 0.1 | 0.4 | n/a | 0.2 | 1.9 | 2.9 | 3.3 | 3.1 | | | | | | | n/a | 4.5 | 3.7 | 5.8 | 6.9 | 5.6 |
| | Neoxaline | | n/a | n/a | n/a | 0.3 | 1.1 | 0.5 | n/a | n/a | n/a | 6.1 | 7.9 | 5.1 | | | | | | | n/a | n/a | n/a | 8.0 | 7.6 | 6.3 |
| | Roquefortine F | | n/a | n/a | n/a | 2.6 | 1.8 | 1.8 | n/a | n/a | n/a | 2.5 | 5.3 | 2.7 | | | | | | | n/a | n/a | n/a | 11.7 | 5.7 | 3.9 |
| | Roquefortine M | | n/a | n/a | n/a | 1.1 | 1.0 | 0.5 | n/a | n/a | n/a | 1.2 | 2.5 | 1.0 | | | | | | | n/a | n/a | n/a | 0.9 | 1.3 | 1.2 |
| | Roquefortine N | | n/a | n/a | n/a | 0.1 | 0.6 | 0.8 | n/a | n/a | n/a | 0.6 | 0.4 | 0.9 | | | | | | | n/a | n/a | n/a | 0.5 | 1.5 | 1.4 |
| Fungisporin-associated | cyclo-(YFVV) | change in peak are, relative to DS27472 | 2.5 | 0.8 | 1.7 | 1.3 | 1.7 | 3.6 | n/a | n/a | n/a | n/a | n/a | n/a | 0.2 | 0.0 | 0.1 | 0.0 | 0.3 | 0.2 | | | | | | |
| | cyclo-(YWVV) | | n/a | n/a | n/a | n/a | n/a | n/a | n/a | n/a | n/a | n/a | n/a | n/a | n/a | n/a | n/a | n/a | n/a | n/a | | | | | | |
| | cyclo-(FFVV) - Fungisporin | | n/a | n/a | n/a | n/a | 1.1 | 0.8 | n/a | n/a | n/a | n/a | n/a | n/a | n/a | n/a | n/a | n/a | 1.3 | 1.8 | | | | | | |
| | cyclo-(YFIV) | | n/a | n/a | n/a | n/a | n/a | n/a | n/a | n/a | n/a | n/a | n/a | n/a | n/a | n/a | n/a | n/a | n/a | n/a | | | | | | |
| | cyclo-(FWVV) | | 0.8 | 0.2 | 0.2 | 1.8 | 1.6 | 3.7 | n/a | n/a | n/a | n/a | n/a | n/a | n/a | 0.1 | 0.9 | 7.6 | 25.1 | 19.0 | | | | | | |
| | cyclo-(FFVI) | | n/a | n/a | n/a | n/a | n/a | n/a | n/a | n/a | n/a | n/a | n/a | n/a | n/a | n/a | n/a | n/a | n/a | n/a | | | | | | |
| | cyclo-(YWVI) | | n/a | n/a | n/a | n/a | n/a | n/a | n/a | n/a | n/a | n/a | n/a | n/a | n/a | n/a | n/a | n/a | 3.3 | 3.2 | | | | | | |
| | YFVV, VYFV, FVVY | | n/a | 2.3 | 1.2 | 1.1 | 0.3 | 0.2 | n/a | 3.5 | 4.7 | 15.1 | 16.9 | 8.0 | n/a | 7.1 | 2.7 | 2.2 | 2.1 | 2.0 | | | | | | |
| | YWVV,VYWV, WVVY | | n/a | 2.4 | 1.1 | 0.9 | 0.4 | 0.7 | n/a | 1.0 | 0.3 | 1.8 | 7.3 | 4.9 | n/a | 7.6 | 1.9 | 2.8 | 8.8 | 3.5 | | | | | | |
| | FVIY, IYFV, FIVY, VYFI | | n/a | n/a | 0.3 | 0.5 | 0.4 | 0.1 | n/a | n/a | 1.4 | 1.9 | 6.2 | 2.9 | n/a | n/a | 0.4 | 2.9 | 8.7 | 1.7 | | | | | | |
| | VFWV, FWVV, WVVF | | 0.6 | 1.4 | 1.7 | 1.0 | 1.0 | 1.1 | 1.1 | 1.5 | 2.5 | 1.7 | 1.6 | 0.3 | 1.8 | 2.0 | 8.7 | 45.8 | 23.7 | 5.7 | | | | | | |
| | FVVF, VFFV, FFVV | | 0.2 | 1.4 | 1.2 | 1.0 | 0.3 | 0.5 | 1.6 | 2.1 | 2.3 | 3.4 | 5.2 | 1.2 | 3.9 | 4.6 | 2.6 | 3.7 | 6.2 | 3.5 | | | | | | |
| | FVIF, FIVF | | n/a | n/a | 1.1 | 0.4 | 0.7 | 0.2 | n/a | n/a | 2.2 | 1.3 | 1.2 | 0.3 | n/a | n/a | 10.3 | 12.5 | 10.8 | 2.5 | | | | | | |

decreased — increased

peak area relative to DS27472

**Supplementary Information SI6:** Averaged changes of normalized peak areas for molecules associated with chrysogine, roquefortine and fungisporin biosynthesis in strains with removed BGCs relative to the parental strain DS27472. Cultivations were performed in SMP medium using shake flasks and sampling took place in 24-hour intervals starting at day 2 (n= 3 independent shake flasks). All analyzed metabolites had a coefficient of variation below 15% for all time points shown here. n/a = not applicable due to absence in DS27472.

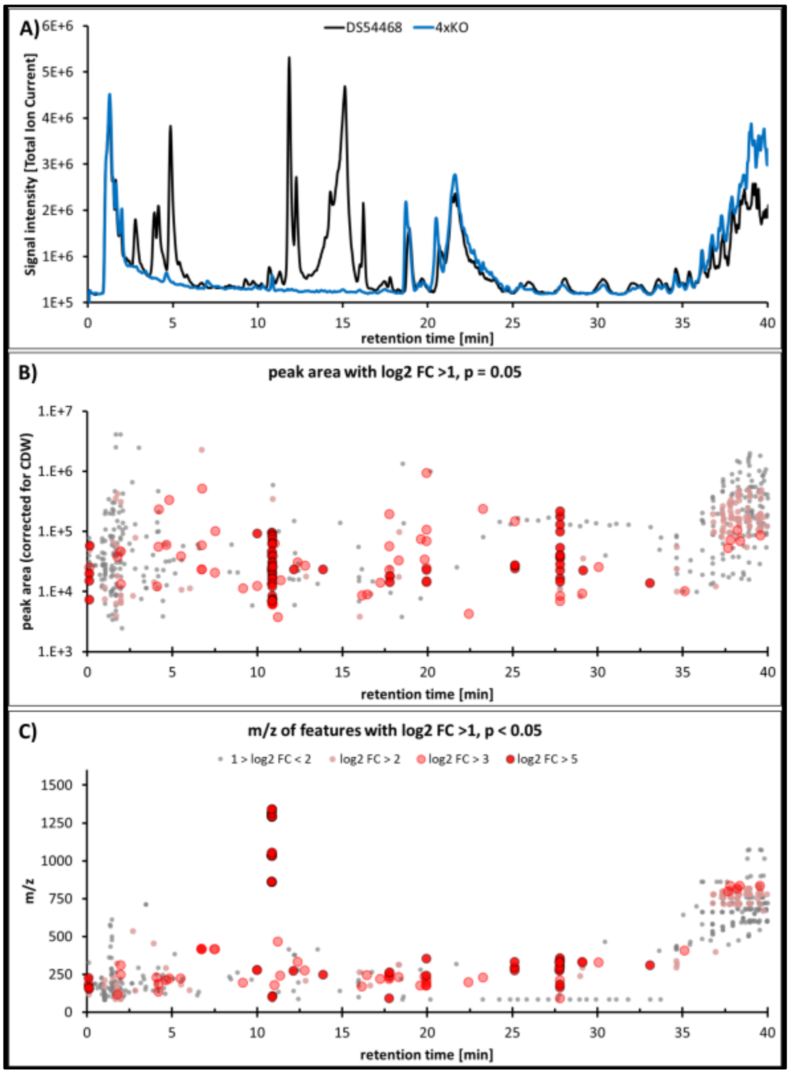


**Supplementary Information SI7:** Comparison of untargeted LC-MS data from shake – flask derived culture broth between DS54468 and 4xKO **A)** Total ion chromatogram (positive ionization mode) showing differences in peaks between DS54468 (black line) and 4xKO (blue line). **B)** Peak area of features enriched in broth samples of 4xKO as identified by XCMS plotted over the retention time. All features with a log2 FC > 1 and p < 0.05 are shown. Each dot represents an individual m/z feature. Color intensity increases stepwise with log2 FC **C)** m/z of features from **B)** plotted over the retention time. Grey dots: 1> log2 FC < 2; small faint red dots: log2 FC > 2; faint red dots: log2 FC >3; bright red dots: log2 FC > 5.

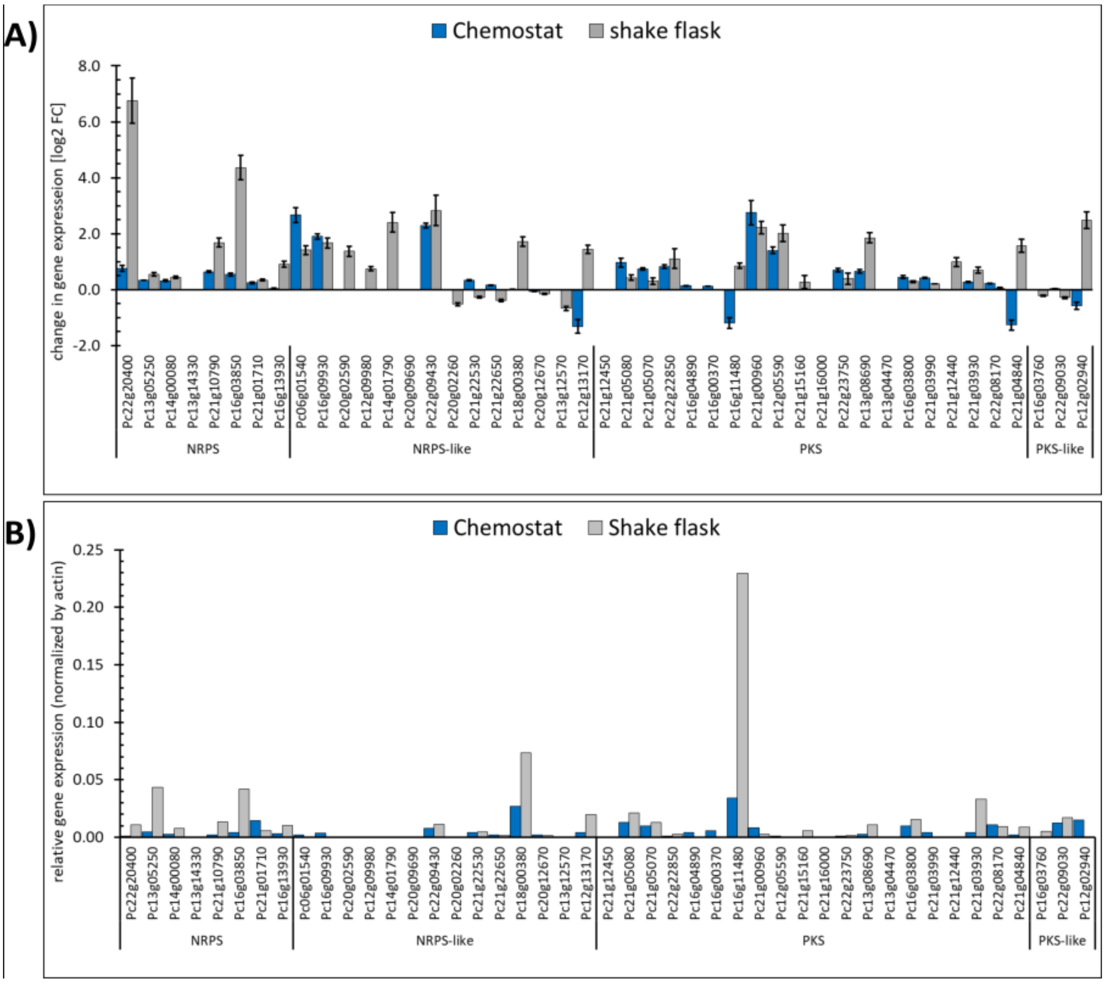


**Supplementary Information SI8:** Transcription changes of remaining BGC core genes in 4xKO compared to DS54468. A) Barplot of log2 FC of SM core gene expression in 4xKO compared to DS54468 in chemostat (blue bars) and shake flask (grey bars) cultures. Bars are missing when genes were not expressed and remained silent. B) Relative gene expression of SM core genes in 4xKO grown in chemostat and shake-flask cultures. Expression is normalized to expression level of actin. Bars are missing when genes are not expressed and remained silent.

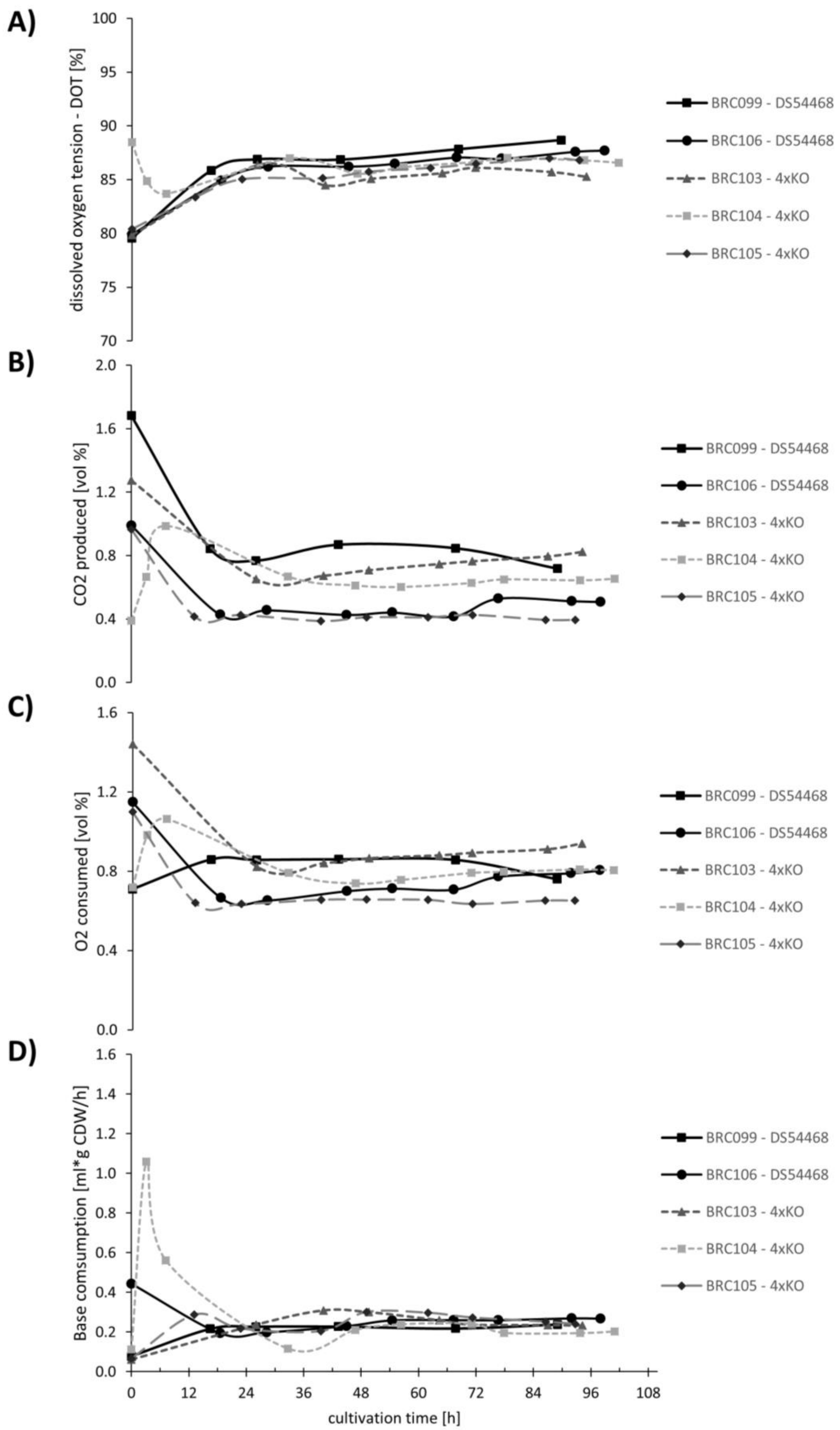
A)
dissolved oxygen tension - DOT [%]
100
95
90
85
80
75
70
BRC099 - DS54468
BRC106 - DS54468
BRC103 - 4xKO
BRC104 - 4xKO
BRC105 - 4xKO
B)
CO2 produced [vol %]
2.0
1.6
1.2
0.8
0.4
0.0
BRC099 - DS54468
BRC106 - DS54468
BRC103 - 4xKO
BRC104 - 4xKO
BRC105 - 4xKO
C)
O2 consumed [vol %]
1.6
1.2
0.8
0.4
0.0
BRC099 - DS54468
BRC106 - DS54468
BRC103 - 4xKO
BRC104 - 4xKO
BRC105 - 4xKO
D)
Base comsumption [ml*g CDW/h]
1.6
1.4
1.2
1.0
0.8
0.6
0.4
0.2
0.0
BRC099 - DS54468
BRC106 - DS54468
BRC103 - 4xKO
BRC104 - 4xKO
BRC105 - 4xKO
0
12
24
36
48
60
72
84
96
108
cultivation time [h]

**Supplementary Information SI9:** Bioreactor-Data of performed cultivations during chemostat phase of bioreactor at a dilution rate of D = 0.05 $h^{-1}$. **A)** Dissolved oxygen tension (DOT). **B)** Produced carbon dioxide **C)** Consumed oxygen. **D)** Base (NaOH) consumption in ml per hour, normalized by cell dry weight (CDW). All parameters did not significantly differ (p>0.05, 2-tailed students t-test) between cultivations when data points after 60 h cultivation were compared between DS54468 and 4xKO.

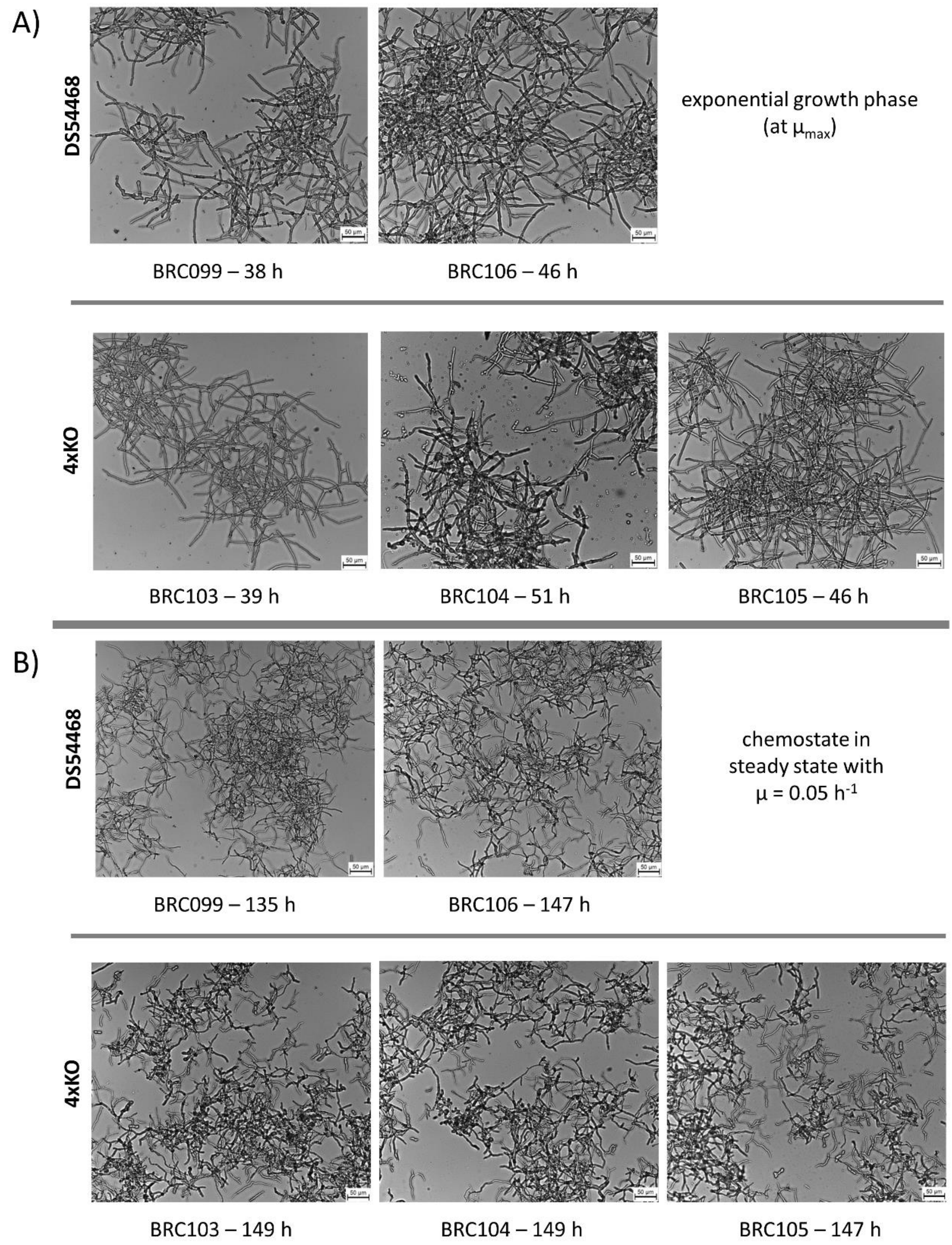

**Supplementary Information SI10:** Light microscopy of mycelium samples of DS54468 and 4xKO. Images are representative for the culture during **A)** exponential and **B)** steady-state growth phase. For glucose-limited chemostats, images were taken after four residence times at a growth rate of 0.05 $h^{-1}$. Scale bar = 50 µm.

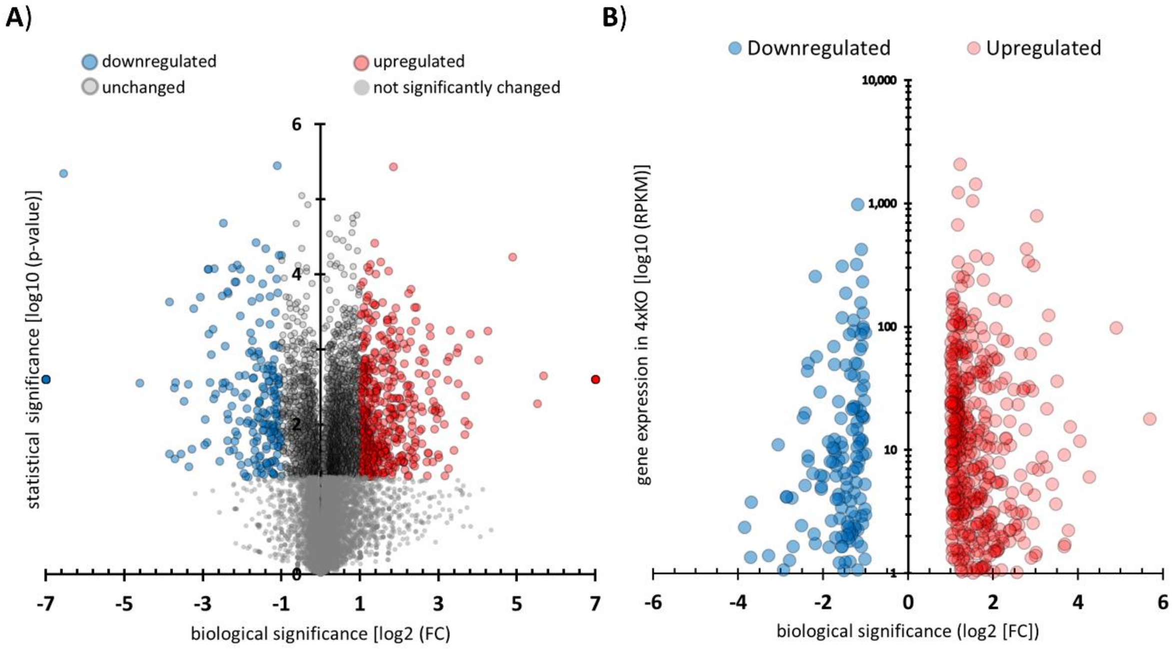


**Supplementary Information SI12**: Visualization of transcriptome changes in 4xKO compared to DS54468 when grown in glucose-limited chemostat at a dilution rate of 0.05 $h^{-1}$. A) Volcano plot displaying statistical significance as log10 (p-value) over biological significance log2 (FC). B) Distribution of expression intensity of up- and downregulated genes in 4xKO. Expression intensity of genes with no statistical significance are not shown. The expression in 4xKO as log10 (RPKM) is plotted on the y-axis over the biological significance log2 (FC) on the x-axis.

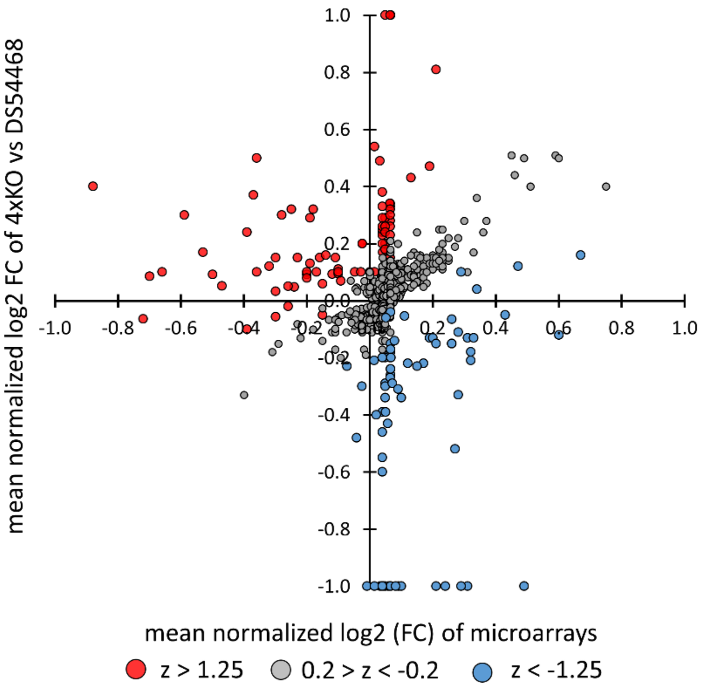


**Supplementary Information SI14.** Scatter plot of mean normalized log2 Fold changes (log2 FC) of RNA-seq results from this study against the mean log2 FC of microarray data distribution. Z-scores above 0.2 and below -0.2 do indicate a weaker correlation between log2 FC observed between the 2 conditions, whereas z-scores closer to 0 do reflect a higher correlation.

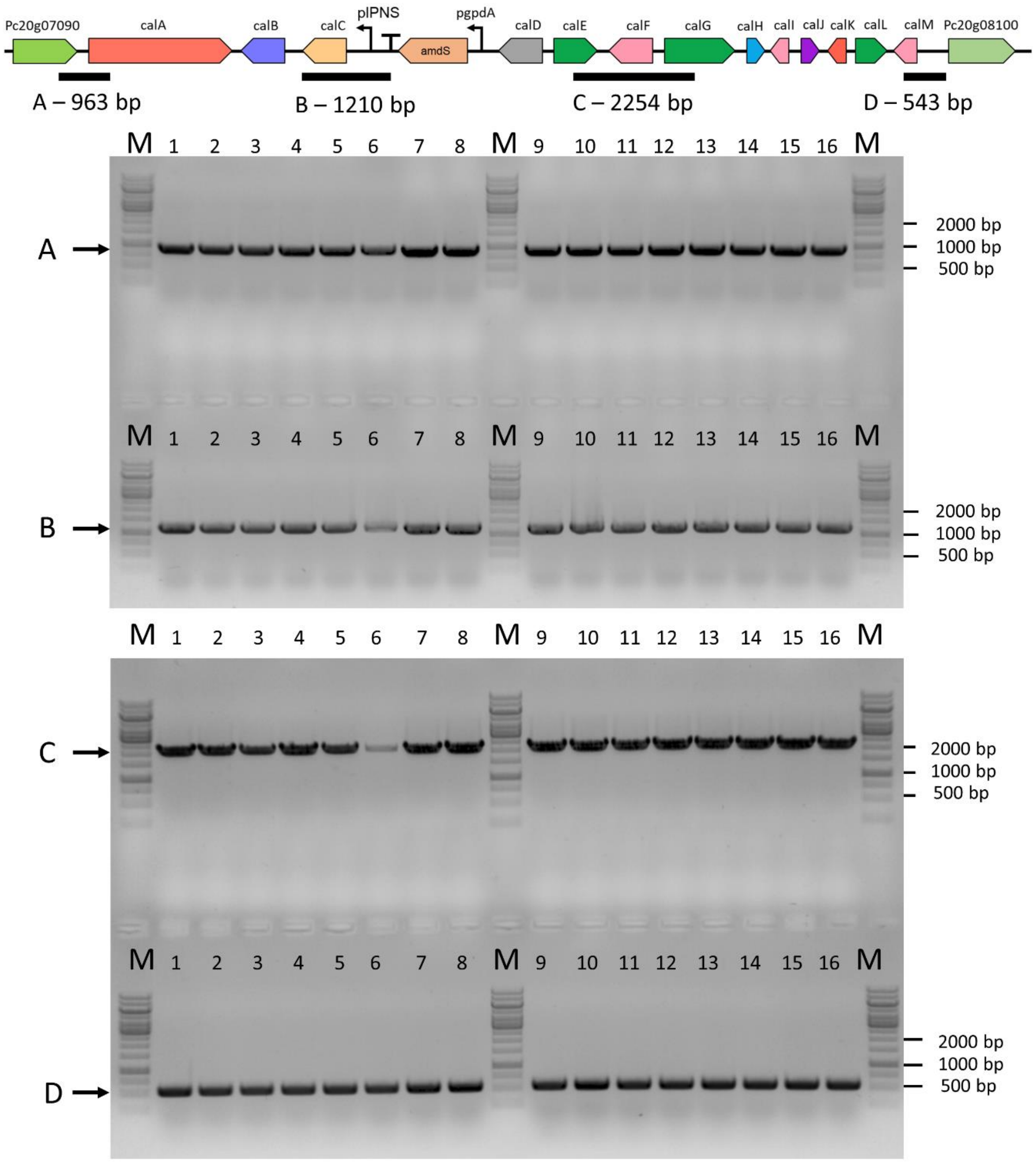


**Supplementary Information SI17**: Colony PCR verification of integration of calbistrin Cluster into 4xKO-B. 16 clones were randomly selected from transformation plates and screened with four primer combinations for presence and correct recombination of transformed DNA parts. PCR amplified regions and expected size of amplicons are depicted on top.

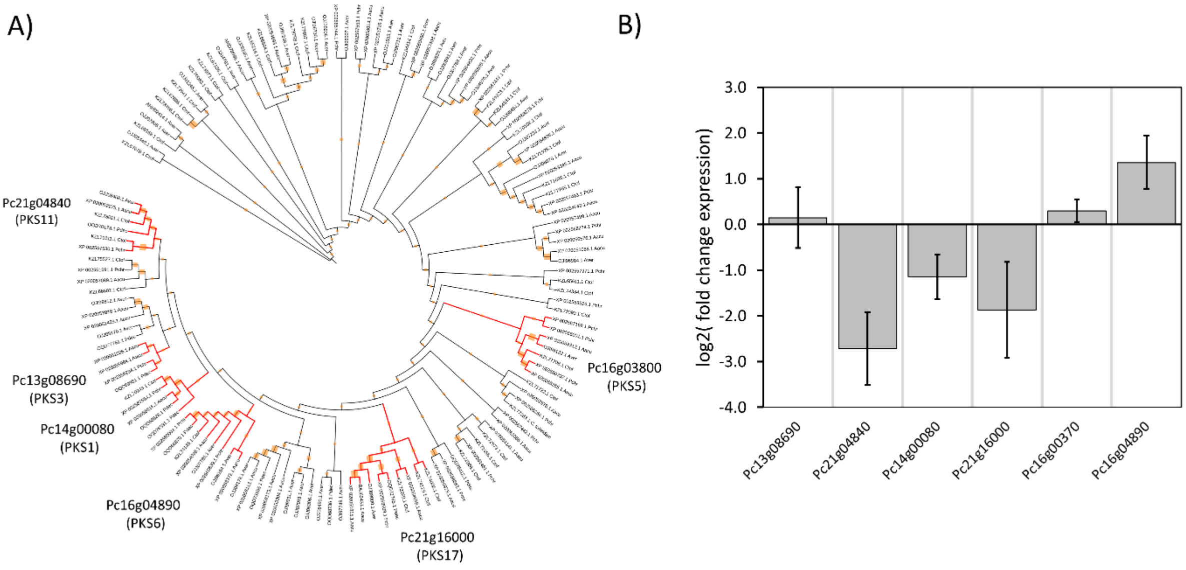


**Supplementary Information SI19:** A) Bootstrap consensus tree of PKS proteins from *P. decumbens*, *A. versicolor*, *A. aculeatus*, *C. tofieldiae* and *P. rubens*. Clustering of proteins from all strains was considered as an indicator that the PKS is involved in formation of the linear moiety of calbistrin. Clustered PKS that were examined are colored in red on the radial tree. B) expression changes of PKS in 4xKO-B-CalBGC strains (n=3) on day 5 of growth in SMP medium.

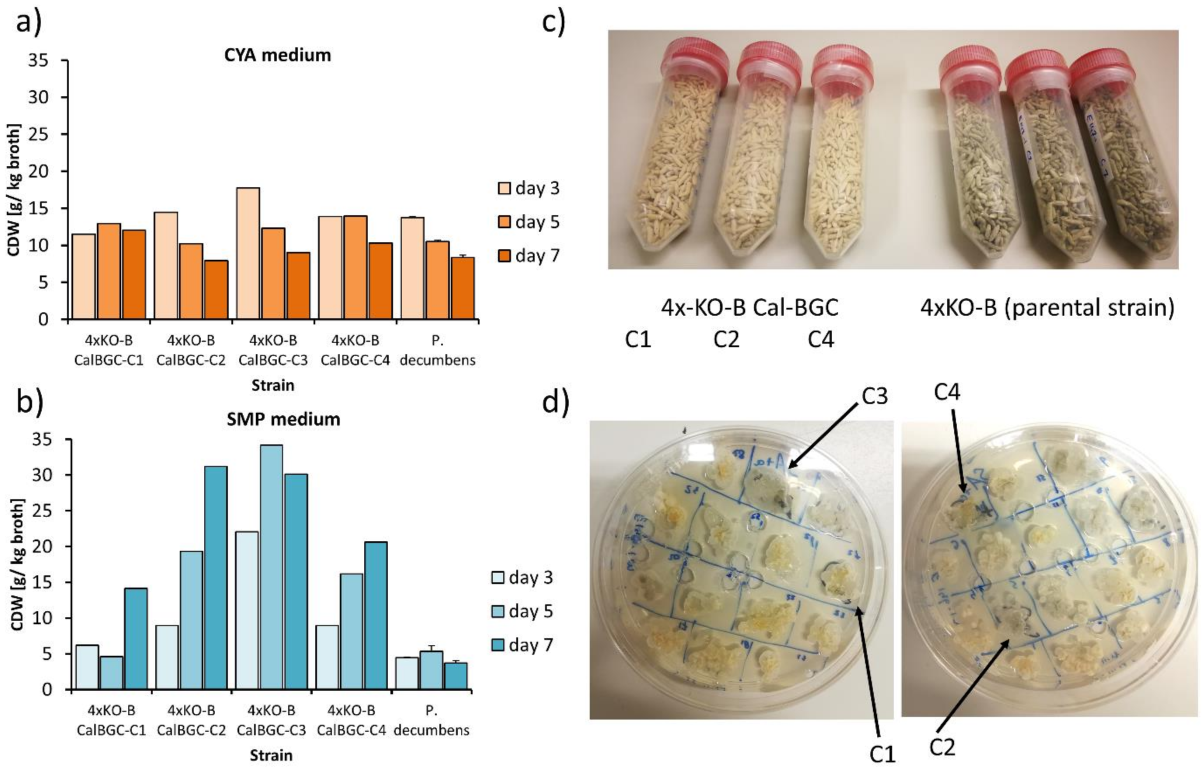


**Supplementary Information SI20**: Impact of calbistrin cluster transformation on growth of *P. rubens* 4xKO-B. A) and B): Biomass formation of four individual clones (CalBGC-C1 to -C4) transformed with the calbistrin BGC. Pure strains were cultivated in shake flasks using CYA and SMP medium. *P. decumbens* was included as a control. Clone CalBG-C3 is a non-producer of decumbenone and thus shows higher accumulation of biomass. C) Appearance of rice batches of CalBGC-transformed clones and the parental strain 4xKO-B. Pigmentation of spores is heavily affected. D) Growth of initially picked transformants on SMP-Agar. Several colonies display a lack of the yellow color associated with production of decumbenones. All clones were verified previously as positive based on colony PCR. Arrows indicate clones that were used for production of calbistrins and decumbenones.
